# Acute changes in systemic glycaemia gate access and action of GLP-1R agonist on brain structures controlling energy homeostasis

**DOI:** 10.1101/2020.07.11.198341

**Authors:** Wineke Bakker, Casper Gravesen Salinas, Monica Imbernon, Daniela Herrera Moro Chao, Rim Hassouna, Chloe Morel, Claire Martin, Giuseppe Gangarossa, Raphael GP Denis, Julien Castel, Andreas Peter, Martin Heni, Walter Maetzler, Heidi Solvang Nielsen, Manon Duquenne, Anna Secher, Jacob Hecksher-Sørensen, Thomas Åskov Pedersen, Vincent Prevot, Serge Luquet

**Author notes:** Corresponding author: Serge Luquet.

## Abstract

The control of body weight and glucose homeostasis are the bedrock of type 2 diabetes medication. Therapies based on co-administration of glucagon-like peptide-1 (GLP-1) long-acting analogues and insulin are becoming popular in the treatment of T2D. Both insulin and GLP-1 receptors (InsR and GLP1-R, respectively) are expressed in brain regions critically involved in the regulation of energy homeostasis, suggesting a possible cooperative action. However, the mechanisms underlying the synergistic action of insulin and GLP-1R agonists on body weight loss and glucose homeostasis remain largely under-investigated. In this study, we provide evidence that peripheral insulin administration modulates the action of GLP-1R agonists onto fatty acids oxidation. Taking advantage of fluorescently labeled insulin and GLP-1R agonists, we found that glucoprivic condition, either achieved by insulin or by 2-deoxyglucose (2-DG), acts as a permissive signal on the blood-brain barrier (BBB) at circumventricular organs, including the median eminence (ME) and the area postrema (AP), enhancing the passage and action of GLP-1-R agonists. Mechanistically, this phenomenon relied on the release of tanycyctic vascular endothelial growth factor A (VEGF-A) and it was selectively impaired after calorie-rich diet exposure. Finally, we found that in human subjects, low blood glucose also correlates with enhanced blood-to-brain passage of insulin suggesting that changes in glycaemia also affect passage of peptide hormones into the brain in humans.

In conclusion, we describe a yet unappreciated mechanism by which acute variations of glycaemia gate the entry and action of circulating energy-related signals in the brain. This phenomenon has physiological and clinical relevance implying that glycemic control is critical to harnessing the full benefit of GLP-1R agonist co-treatment in body weight loss therapy.

## INTRODUCTION

Obesity and correlated diseases are now clearly identified as a worldwide pandemic in both developing and developed countries (Molavi et al., 2006) and have more than doubled since 1980 (World Health Organization, WHO 2015). Obesity is associated with increased mortality, notably due to a constellation of associated disorders such as type 2 diabetes (T2D), cardiovascular, gastrointestinal, reproductive diseases, non-alcoholic fatty liver disease and certain cancers, thus impacting upon large swathes of society and placing an enormous burden on health care resources. The last decade has witnessed a significant endeavor to efficiently develop alternative strategies aiming at restoring glycaemia control and decreasing body weight.

While insulin treatment remains a major tool in the therapeutic arsenal to restore uncontrolled glycaemia in diabetes, other existing therapies are based on peripheral peptides whose respective receptors are also located in the central nervous system (CNS). Among these peptides, the glucagon-like peptide-1 (GLP-1), which is notably secreted from gut enteroendocrine cells upon nutrient transit, exerts several beneficial actions partly through its ability to enhance glucose-induced insulin release from pancreatic beta-cells (Barrera et al., 2011; Muller et al., 2019). This incretin action has fostered the development of pharmacological agonists of GLP-1 receptor (GLP-1R) for the treatment of T2D and obesity. Pharmacological interventions combining insulin and GLP-1R agonists are on the market for the treatment of T2D, yet limited knowledge exists on the mechanistic interactions of these two molecules (Anderson and Trujillo, 2016; Gough et al., 2014; Moreira et al., 2018). Importantly, in addition to the incretin effects (Anderson and Trujillo, 2016), GLP-1 promote satiety and lowers body weight (Muller et al., 2019). These actions are believed to be, at least in part, mediated through brain circuits dedicated to the regulation of energy balance (Dodd and Tiganis, 2017; Taouis and Torres-Aleman, 2019).

Peripheral insulin is reported to access the brain either through regulated passage across the blood brain barrier (BBB) (Banks et al., 1997; Pardridge et al., 1985) or via circumventricular organs (CVOs) (Banks, 2019) and locally initiates signaling cascades through its cognate insulin receptor (IR), which is highly expressed in several hypothalamic and extra-hypothalamic regions (Konner et al., 2011; Vogt and Bruning, 2013). In both humans and rodents, impaired central insulin signaling has been associated with metabolic defects (Bruning et al., 2000; Dodd and Tiganis, 2017; Ferrario and Reagan, 2018; Fisher et al., 2005; Heni et al., 2015; Scherer et al., 2011).

Peripheral GLP-1 has also been implicated in body weight homeostasis. GLP-1 was found to modulate vagal afferents, while centrally produced GLP-1 can also serve as a central neurotransmitter (Burcelin and Gourdy, 2017; Lockie, 2013) (Baggio and Drucker, 2014). At the central level, GLP-1 is produced by a distinct subset of neurons within the nucleus tractus solitarius (NTS) and several studies now point towards an unequivocal role of GLP-1-dependent signaling in the control of body weight, food intake and energy expenditure. Recent studies suggest that peripherally administered GLP-1R agonists act on GLP-1R-expressing neuronal populations located in, and in the vicinity of, the circumventricular organs (CVOs), such as the anorectic/catabolic pro-opiomelanocortin (POMC) neurons and/or the orexigenic/anabolic agouti-related peptide (AgRP) neurons of the arcuate nucleus (ARC) (Baggio and Drucker, 2014; Secher et al., 2014). Interestingly, while several studies have used direct brain administration of hormones and analogues to dissect out the neural substrates underlying their central actions, there is a paucity of knowledge regarding the physiological mechanisms by which endogenously secreted or pharmacologically administered hormones gain access to effector neurons located behind the BBB. Among the CVOs, the median eminence (ME) and the area postrema (AP), located near the 3rd and 4th ventricles respectively, are gate points for circulating signals acting on the ARC and NTS neurons (O’Rahilly and Farooqi, 2008a, b), two key structures that regulate body weight dynamics. While being characterized by reduced fenestrated capillaries when compared to the ME, the ARC is enriched in tanycytes, a specialized glial cell-type (Garcia-Caceres et al., 2019; Prevot et al., 2018). The tanycytes are believed to control the fenestration of blood vessels at the ME/ARC junction, and facilitate transport of blood circulating signals onto energy sensing ARC neurons (Balland et al., 2014; Langlet et al., 2013a; Mullier et al., 2010; Rodriguez et al., 2010).

In particular, tanycytes have been shown to gate adaptive remodeling within the ME by controlling blood-brain signal exchanges in response to nutrients availability and plasma glucose (Langlet et al., 2013a) (Balland et al., 2014). We speculated that acute physiological fluctuations (plasma glucose, nutrients or hormones) might regulate the ability of hormones and/or pharmacological analogues to exert their central actions by modulating their access to central structures. Considering the growing therapeutic need of poly-pharmacological strategies relying e.g. on the co-administration of GLP-1R agonists and insulin (Anderson and Trujillo, 2016), we seek to explore whether and how insulin and GLP-1R analogues might exert coordinated/synergistic actions on energy homeostasis/physiology through direct and/or indirect modulation of BBB passage.

Using bioactive fluorescently-labelled insulin and the GLP-1R agonist Exendin-4 (Ex-4), we show that peripherally administered insulin can initiate signaling cascades in the ME/ARC and potentiate accelerated Ex-4 access to AP/NTS and ME/ARC structures. In addition, we show that central detection of hypoglycemia, rather than direct insulin action, is instrumental to enhance Ex-4 brain access. While GLP-1R agonist injection alone induced a sustained increase in fatty acid oxidation, upon co-injection with insulin, Ex-4 was able to partially oppose the lipogenic effect of insulin. We provide evidence that the release of vascular endothelial growth factor A (VEGF-A) by tanycyte relay the adaptive response of CVO to glycemic change.

Finally, we show that exposure to an energy-rich diet leads to a specific uncoupling between hypoglycemia and the potentiation of Ex-4 entry in the brain.

Altogether, our data reveal that acute changes in systemic glycaemia can rapidly modulate the access and action of peripheral metabolic signals onto relevant energy-related neurons behind the BBB, in regions that contain or are in close vicinity to tanycytes. Finally, in support of this hypothesis, we demonstrate that peripheral blood glucose negatively correlates with the ratio of central cerebrospinal fluid (CSF) to systemic plasma insulin in humans, further indicating that low plasma glucose may be associated with increased blood-to-brain passage of metabolically active peptide hormones. The identified mechanism is of clinical relevance as combined therapy used for glycemic control in diabetes could serve as gate-keeper for brain access and action of GLP1/insulin crucial for whole-body energy homeostasis.

## RESULTS

### Co-administration of insulin and Exendin-4 reveals bidirectional metabolic actions

We first explored the metabolic consequences of insulin and Ex-4 co-administration. In three groups of male C57Bl6J mice (n=8/group) we measured the metabolic efficiency using indirect calorimetry, feeding patterns and locomotor activity during a 4-days treatment period consisting of daily injections (2:00 pm) of insulin (20 nmol/kg, blue), Ex-4 (120 nmol/kg, red) or a mix of insulin+Ex-4 (20 nmol/kg, 120 nmol/kg, green) (**Figure 1A**). A 3-days baseline period was first acquired for all groups (n=24) and presented as reference (black, **Figure 1A**). Consistent with previously described anorectic actions of GLP-1 (Baggio and Drucker, 2014; Barrera et al., 2011; Burcelin and Gourdy, 2017; Secher et al., 2014; Sisley et al., 2014), acute administration of Ex-4 decreased both daily (**Figure 1B**) and cumulative (**Figure 1C**) food intake either alone or in combination with insulin. These satiety-like responses were associated with reduced energy expenditure when compared to control group (**Figure 1D**). Consistent with the body weight loss effect induced by the GLP-1R agonist, acute injection of Ex-4 triggered a sustained increase in fatty acids oxidation compared to control baseline (**Figure 1E**), and both Ex-4 and Ex-4+insulin treatments were associated with a significant decrease in body weight compared to control condition (**Figure 1G**). Analysis of meal ultrastructure revealed that Ex-4 was efficient in decreasing meal number and meal size following its injection, even though a desensitization processes may occur during the 4-days treatment (**Figure S1**). Consistent with feeding response to hypoglycaemia (Fraley and Ritter, 2003; Hudson and Ritter, 2004), acute insulin administration led to a transient increase in food intake (**Figure 1B, C**). This response was associated with a slight decrease in nocturnal chow intake and overall cumulated food intake was significantly different from control only during the first day (**Figure 1C**). It is worth to note that, while energy expenditure remained unaffected when compared to control (**Figure 1D**), acute insulin administration exerted a sharp and sustained decrease in fatty acids oxidation (**Figure 1E**) consistent with the anti-lipolytic and lipogenic action of insulin (Petersen et al., 1988; Thomas et al., 1979). Strikingly, when co-administered with insulin, Ex-4 opposed the hyperphagic response following hypoglycaemia (**Figure 1B**) and induced a sharp rebound in fatty acids oxidation (**Figure 1E**). This result reveals antagonistic actions of GLP-1R agonist on feeding and metabolic substrates utilization, even under co-administration with insulin. Note that we only observed a decrease in locomotor activity when insulin and Ex-4 were co-administered (**Figure 1F**) as a possible result of reduced energy availability and/or low blood glucose levels. Our results support the idea that body weight loss achieved by Exendin-4 is due to both reduction of feeding and shift toward peripheral lipids utilization (Sisley et al., 2014). We next explored whether the effect was a consequence of a peripheral or brain-specific action.

**Figure 1.**
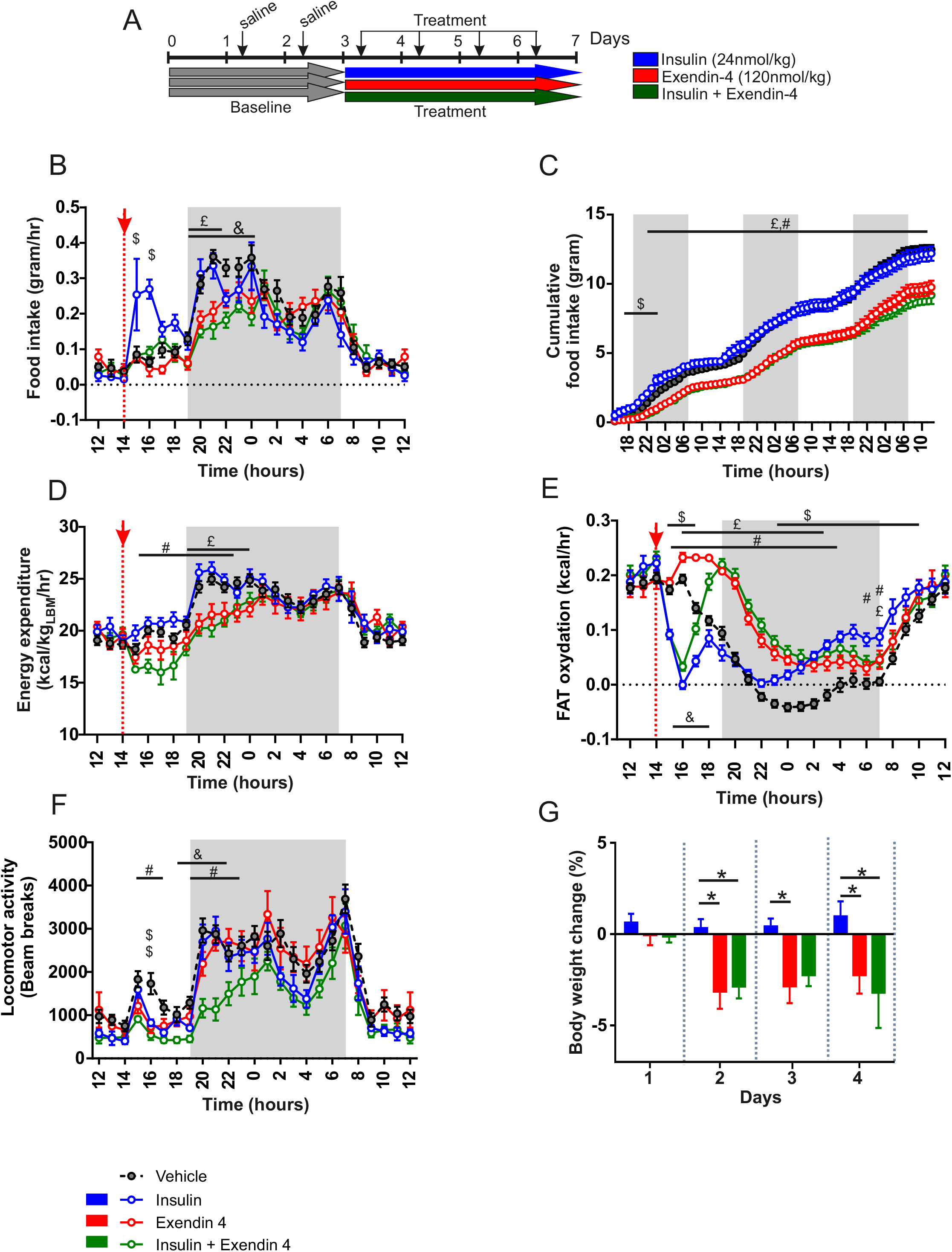
Metabolic consequences of insulin and Exendin-4 co-injection. (**A**) Experimental set up for metabolic efficiency characterization of mice receiving a daily i.p. saline injection (baseline, grey) followed by a 4-days treatment period consisting of a daily injection (2:00 pm) of insulin (20 nmol/kg, blue), Exendin-4 (120 nmol/kg, red) or mix of insulin+Exendin-4 (20 nmol/kg, 120 nmol/kg, green). Graphs represent an average of a 3-days baseline (grey) or treatment periods 3-days treatment for (**B**) food intake, (**C**) cumulative food intake, (**D**) energy expenditure, (**E**) calculated fatty acids oxidation, (**F**) locomotor activity. (**G**) Body weigh changes upon treatment. N=8 in each group. Data are expressed as mean ± SEM. **\*** indicated P value<0.05. $ indicated P value<0.05, insulin vs vehicle. £: indicated P value<0.05, Ex-4 vs vehicle. #: indicated P value<0.05, Insulin+Ex-4 vs vehicle. &: indicated P value<0.05, Insulin+Ex-4 vs Ex-4.

### Peripheral insulin accesses hypothalamic CVOs initiating IR-dependent signaling cascade

Using biologically active, fluorescently labelled insulins (vivotag-750 labelled insulin: insulin_VT^750^ or Alexa^647^ labelled insulin: insulin_Alexa^647^) in combination with whole brain light sheet fluorescence microscopy (LSFM) technology (Salinas et al., 2018), we assessed whether insulin could directly access the brain. We found that peripherally administered fluorescent insulin can indeed access the ME and ARC hypothalamic regions (**Figure S2A-F**). Interestingly, this signal required an IR-dependent mechanism since mice pre-treated with the IR antagonist S961 (1060 μmol/kg) 30 min prior systemic administration of insulin_VT^750^ failed to show fluorescent signals in the ARC (**Figure S2A, B**). To confirm our findings, the same experiment was repeated with Insulin-Alexa^647^ using confocal microscopy. Mice administered with Insulin-Alexa^647^ showed similar signal patterns (**Figure S2C, D**). Co-labelling analysis revealed that, 2 min after intravenous (iv) injection of Insulin-Alexa^647^, fluorescent insulin signals were detected in ME cellular elements expressing vimentin, a specific marker of tanycytes (**Figure S2E, F**). As with insulin_VT^750^, insulin_Alexa^647^ was almost completely absent in the ARC when mice were pre-treated with S961 prior to the administration (**Figure S2E, F**). In addition to the entry of insulin in the brain, immunofluorescence analysis revealed that, after 10 min of iv injection, insulin was able to trigger IR downstream phosphorylation of Akt (Ser^473^) in ME/ARC regions with an intense signal in vimentin-expressing tanycytes (**Figure S2G**). Western blot analysis of ME/ARC punches, compared to whole mediobasal hypothalamic (MBH) punches, revealed that phospho-Akt (ser^473^), but not phospho-ERK (Thr^202^/Thr^204^), was preferentially induced in ME/ARC by peripheral insulin injection (**Figure S2H-J**).

### Insulin and GLP-1 signaling components are differentially expressed in hypothalamic and hindbrain tanycytes

Using similar approaches, we found that peripheral administration of Ex-4_Alexa^594^ was detected in ME/ARC and AP (**Figure 2A-D**) structures. Immediately after Ex-4 IV injection (10-15 sec), the fluorescent signal was observed in both ME and AP (**Figure 2A, B**) however, a clear accumulation of signal was visible in the ME tanycytes (**Figure 2A, B**) suggesting that in early time point only Exendin-4 transport involved ME tanycytes while diffusing through fenestrated capillaries in the AP. In addition, 15 min after fluorescent Ex-4 IP injection, the signal was detected in the ARC and overlapped with GLP-1R immunoreactivity (**Figure 2C, D**).

**Figure 2.**
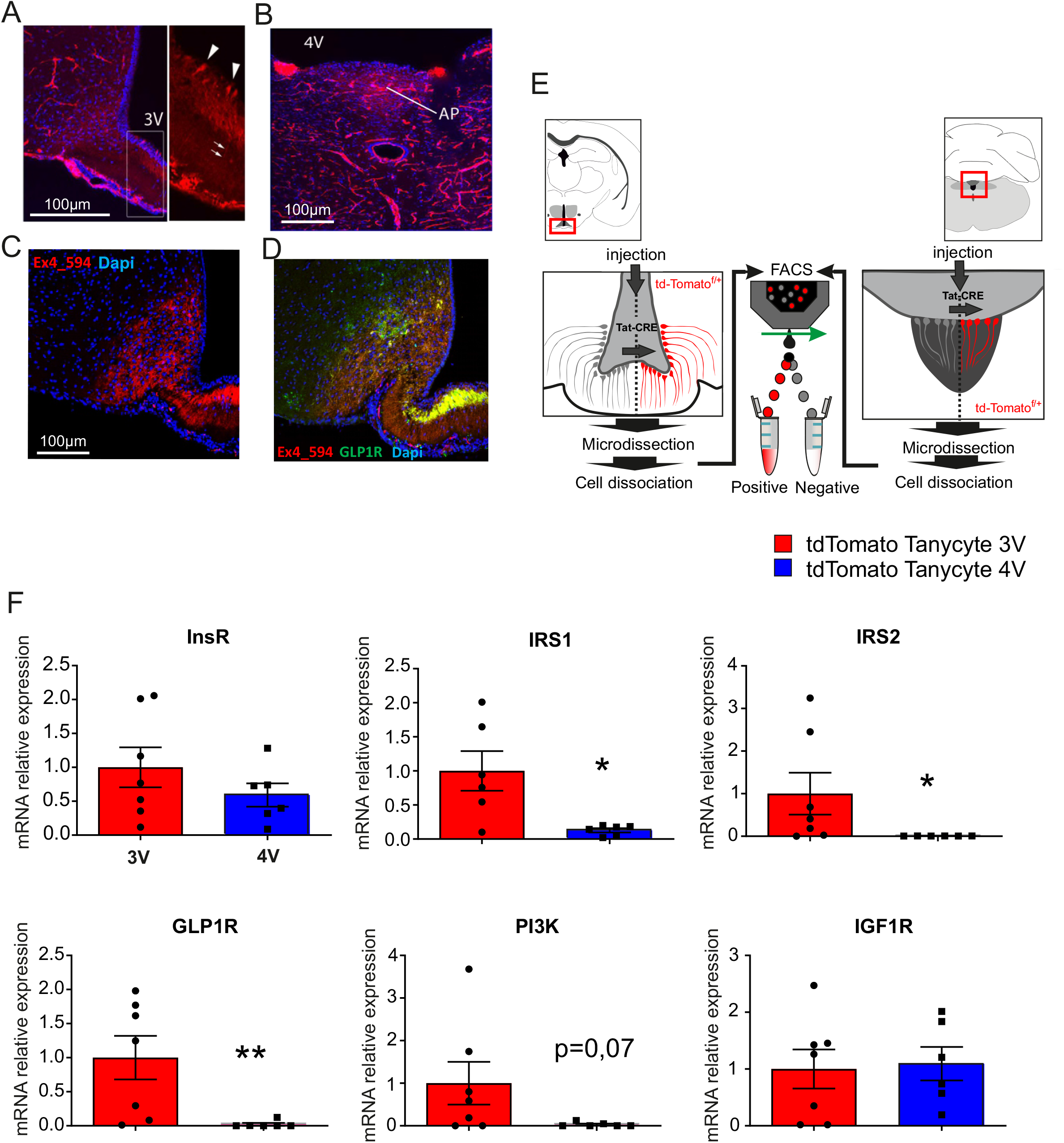
GLP-1R and Insulin-R expression and function in ME/ARC and AP/NTS region. (**A-D**) Representative photomicrographs of ME/ARC (**A, C, D**) and AP (**B**) of fluorescent Exendin-4_Alexa^594^ (**A, C**) and counterstained with DAPI (**B**) or immunochemical detection of GLP-1 R (**D**). (**A, B**) animals were sacrificed 10-15 sec after peripherally injected fluorescent Exendin-4_Alexa^594^ while photos in **C, D** correspond to animals sacrificed 15 min after Exendin-4_Alexa^594^ injection. (**E**) Protocol for tanycyte FACS isolation and (**F**) relative mRNA content for Ins-R and GLP-1R associated machinery in FACS-sorted tanycytes from the ARC/ME (red) or AP/NTS (blue) of tdTomato^loxP/+^ reporter mice following 3rd or 4th ventricle injection of Tat-Cre protein.

To understand the potential differences between ME and AP tanycytes, we performed FACS purification of fluorescently labelled tanycytes (Langlet et al., 2013a). Briefly, Tat-Cre chimeric protein was injected either in the 3rd or 4th ventricle of tdTomato^loxP/+^ mice, thus allowing cell-type specific tagging of tanycytes (Langlet et al., 2013a). Labeled tanycytes were then subjected to FACS sorting (**Figure 2E**) followed by RT-PCR transcript analysis. As shown in **Figure 2F**, transcripts related with signaling pathway of both GLP1, like GLP1 receptor (GLP1R), and insulin, like insulin receptor substrate 1, 2 (IRS1, IRS2) and phosphatidyl inositol 3 kinase (PI3K) were significantly enriched in the tanycytes of the 3rd ventricle (ME) compared to 4th V (AP) even though both ME and AP tanycytes expressed similar amount of mRNA encoding for insulin receptor (InsR) and insulin-like growth factor receptor (IGF1-R transcripts (**Figure 2F**). These data support the idea that only tanycytes from the third ventricle are actively involved in both insulin and exendin-4 transport into the CSF.

### Peripheral insulin administration promotes the access of Exendin-4 into hypothalamic/hindbrain structures

We next explored the action of peripherally injected insulin on brain access of Exendin-4. LSFM-based whole brain analysis (Salinas et al., 2018) was used to quantify the accumulation of Ex-4_VT^750^ after systemic administration. 2D planes extracted from LSFM clearly showed that an Ex-4_VT^750^ signal was detected in discrete brain structures including the ME/ARC, OLVT, AP, SFO and choroid plexus (**Figure 3A**). Compared to vehicle, peripheral injection of insulin (240 nmol/kg) increased the signal of Ex-4_VT^750^ in these brain areas (yellow arrow, **Figure 3A**). Moreover, using 3D projections we could demonstrate that peripheral injection drastically increased Ex-4_VT^750^ fluorescent signal in brain vasculature as well as in ME/ARC regions (**Figure S3**). We next applied a recently developed approach that allows region-specific quantifications of fluorescent signals in an integrated brain atlas (Salinas et al., 2018). Using this method, we found that insulin exerted a dose-dependent action onto Ex-4_VT^750^ hypothalamic access which could be prevented by pre-treatment with the IR antagonist S961 (**Figure 3B**). These results indicate that peripheral insulin injection leads, directly or indirectly, to enhanced passage of Exendin-4 into brain regions were GLP1-R is expressed.

**Figure 3.**
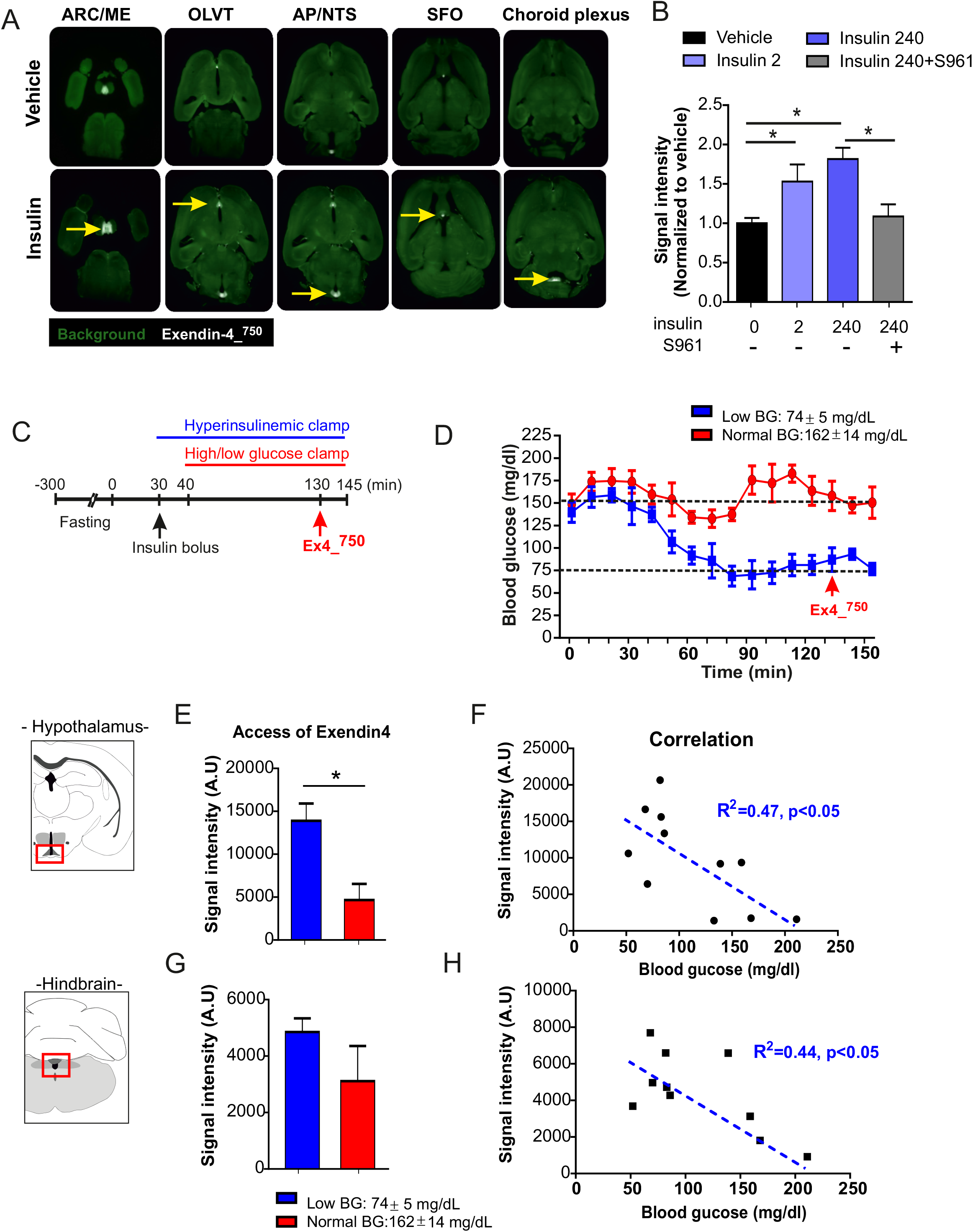
Insulin-mediated hypoglycemia potentiates the access of GLP-1 analogue Exendin-4 into the brain. **(A)** Selection 2D-planes whole brain light sheet scanning to visualize fluorescent signal of peripherally injected Exendin-4_VT^750^ (120 nmol/kg) together with intraperitoneal (ip) injection of vehicle or insulin (240 nmol/kg). The picture represents arcuate nucleus (ARC), organum vasculosum of the lamina terminalis (OVLT), area postrema (AP), subfornical organ (SFO), and choroid plexus while (**B**) Quantification of hypothalamic Exendin-4_VT^750^ fluorescent signal normalized to vehicle in combination with peripheral injection of Insulin (2 nmol/kg, 240 nmol/kg), Insulin (240 nmol/kg) and insulin receptor antagonist S961+insulin. n=5-17 per each group. **\***, indicates P<0.05, vehicle *vs* insulin. Data are expressed as mean ± SEM. (**C**) Experimental set up for hyperinsulinemic clamp controlled at either normal glucose or low glucose levels. (**D**) Low or normal glucose during hyperinsulinemic conditions. (**E, G**) Quantification of Exendin-4_VT750 fluorescent signal normalized to vehicle under high (162±14 mg/dl) or low (74±5 mg/dl) glucose condition in (**E**) hypothalamus and (**G**) hindbrain. (**F, H**) Correlation curve between Exendin-4_VT^750^ fluorescent signal intensity and blood glucose (BG) in (**F**) hypothalamus and (**H**) hindbrain. n=5-9 minimum in each group. **\***, indicated P<0.05, vehicle *vs* insulin. Data are expressed as mean ± SEM.

### Hypoglycemia rather than insulin itself gates central access of GLP-1R agonist

Since the median eminence undergoes adaptive structural plasticity (higher permeability under fasting-induced hypoglycemia (Langlet et al., 2013a)) to match the energy status of the individual, we next explored whether brain insulin signaling *per se* or insulin-promoted hypoglycemia facilitates Ex-4_VT^750^ access in ME and AP. We therefore performed an hyperinsulinemic clamp (3.3 mU/kg/min) on awake mice in which glucose levels were set at either physiological (162±14 mg/ml) or low (74±5 mg/ml) concentrations. Ex-4_VT^750^ was administered intraperitoneally under constant hyperinsulinemia with blood glucose stabilized under normal or low levels (**Figure 3C, D**). Animals were sacrificed 15 min after Ex-4_VT^750^ administration and brains were processed for LSFM-based signal acquisition (Salinas et al., 2018). We found that, in a hyperinsulinemic state, Ex-4_VT^750^ signal was higher in the hypothalamus of animals maintained at low glucose levels compared to the euglycemic controls (**Figure 3E, G**). Furthermore, regression analysis revealed an inverse correlation between low glucose levels and high Ex-4_VT^750^ fluorescence in both hindbrain and hypothalamic structures (**Figure 3F, H**). These data indicate that, although we cannot rule out a direct action of insulin onto different brain regions, changes in glucose levels are necessary to promote Exendin-4 access in the brain.

Next, we explored whether centrally detected glucoprivation could relay CVOs’ adaptation to hypoglycemia. In that regards we assessed whether central neuroglucopenia induced by 2-deoxyglucose (2-DG, 250 mg/kg) peripheral injection could affect brain entry of a GLP1-R agonist. Animals received either saline or 2-DG injections followed by intravenous (?) administration of Ex-4_VT^750^ 30 min after 2-DG injection (**Figure 4A, B**) and ~10 min prior to sacrifice. The brains were dissected and processed as previously indicated 10 minutes after Ex-4_VT^750^. Counterregulatory response to neuroglucopenia was evidenced by increased blood glucose following 2-DG administration (**Figure 4B**). 2-DG injection was associated with an increase in Ex-4_VT^750^ signal in the hindbrain and a trend to increase in the hypothalamus (**Figure 4C, E**). In the hindbrain, we found a significant correlation between blood glucose and Ex-4_VT^750^ signal (**Figure 4F**), indicating that the access of Ex-4_VT^750^ to the brain was associated to the degree of the counterregulatory response. Importantly, while low blood glucose was correlated with an increase in brain passage through CVOs of Ex-4_VT^750^ in our previous observations (**Figure 3, S3 and 4**), here we found that hindbrain Ex-4_VT^750^ fluorescent signal (and to a lesser extend hypothalamus), was increased despite large counterregulatory increase in blood glucose 30 min after 2-DG injection. This observation suggests that the adaptive changes occurring in CVOs to modulate Ex-4 passage might involve the detection of glucoprivation at central level which can potentially be decorrelated from peripheral glucose change.

**Figure 4.**
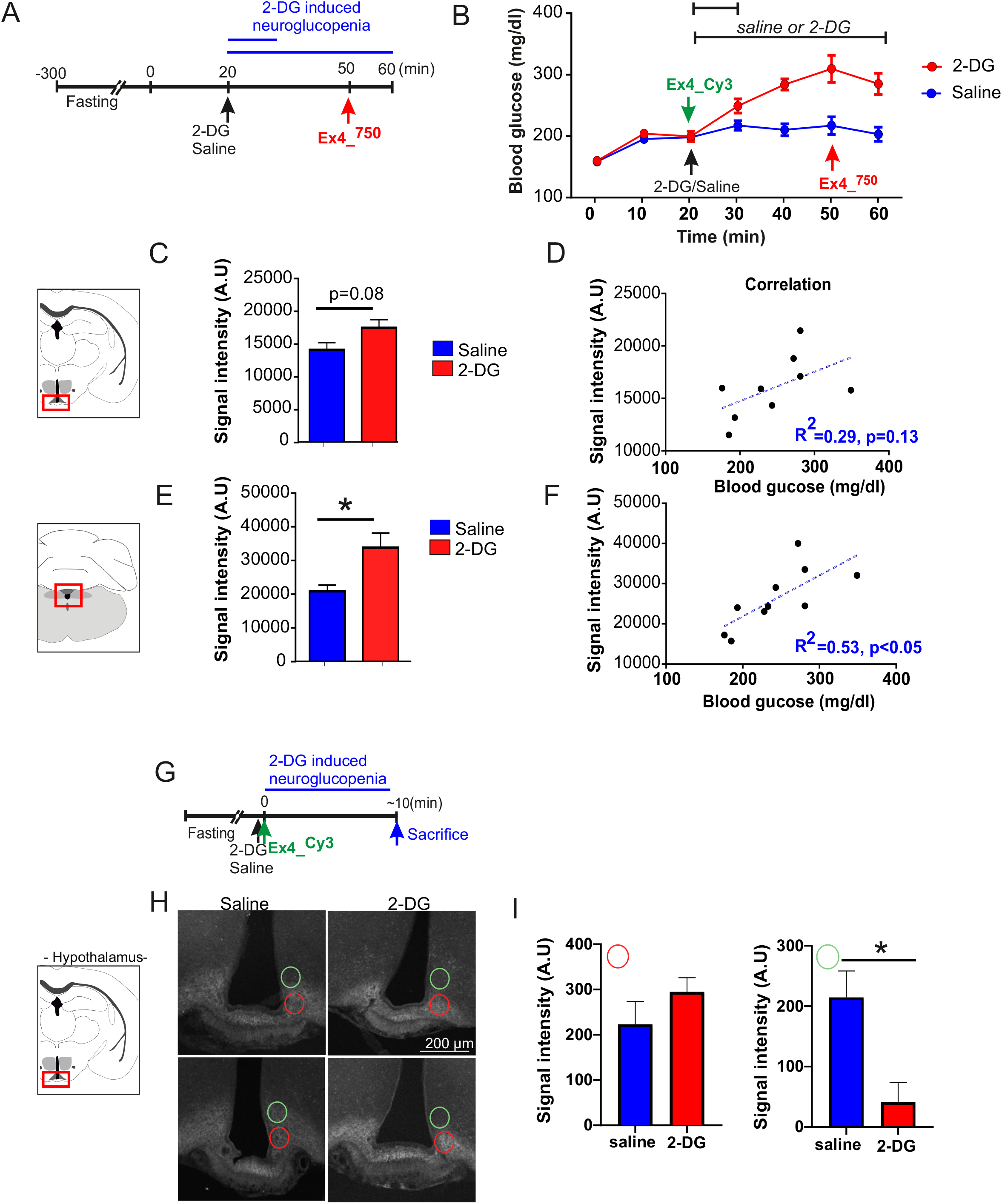
Central neuroglucopenia increases Exendin-4 access to the brain. (**A**) Experimental set up of 2-DG (250 mg/kg) injections prior to peripheral injection of Exendin-4_VT^750^ (120 nmol/kg) at 10 min or 30 min following 2-DG/vehicle injection, (**B**) glycemic change upon 2-DG induced neuroglucopenia. (**C-F**) Quantification of fluorescent signal from Exendin-4_VT^750^ or Exendin-4_^Cy3^ in the hypothalamus and (**C, D**) hindbrain (**E, F**) structure using whole brain laser sheet microscopy. (**D, F**) Correlation curve between Exendin-4_VT^750^ fluorescent signal intensity and blood glucose in (**D**) hypothalamus and (**F**) hindbrain. n=5-9 minimum in each group. **\***, indicated P<0.05. Data are expressed as mean ± SEM.(**G**) Experimental set up for concomitant injection of vehicle or 2-DG (250 mg/kg) together with Exendin-^Cy3^ (120 nMol/kg) 10 min or 30 min following 2-DG/vehicle injection ~15min prioir to sacrifice. (**H**) representative photomicrograph for Exendin-^Cy3^ fluorescent distribution and fluorescent signal quantification (**I**) signal quantification in the upper (green circles) and lower part (red circles) of the arcuate nucleus ~15 min after 2-DG bolus injection. n=5-9 minimum in each group. **\***, indicated P<0.05. Data are expressed as mean ± SEM.

Hence, in order to further precise the dynamic of Exendin-4 entry to the brain following neuroglucopenia, we performed a similar experiment in which animals were sacrificed at early time point (~15min) following the injection of both fluorescent Exendin-4^Cy3^ and 2-DG (**Figure 4 G**) prior to any change in peripheral glucose. Confocal microscopy analysis of brain section in the ARC (**Figure 4 H**) and AP region (**Figure S4**) analysis revealed that even prior to any systemic change in glucose 15 min after 2-DG injection, centrally detected glucoprivation resulted in the redistribution of the fluorescent Exendin-4^Cy3^ more ventrally towards the ME in the ARC regions (**Figure 4 H, I**) while the overall intensity in the ARC and AP remained unchanged (data not shown and **Figure S4**). Overall, these results support the notion that Ex-4 passage through hypothalamic CVOs and its distribution in hypothalamic structures can rapidly respond to change in glycemia detected, at least in part, at central level.

### Tanycytic VEGF contributes to brain access and action of GLP-1 agonist

Vascular endothelia growth factor (VEGF) has been shown to be an important regulator of BBB plasticity (Dantz et al., 2002). In addition, tanycytic VEGF has been involved in the control of ME/ARC adaptations to fasting and hypoglycaemia (Langlet et al., 2013a). We therefore hypothesized that brain access and action of Ex-4 during insulin-mediated hypoglycaemia could depend, at least in part, on tanycytic *Vegfa*. First, to explore the role of VEGF signaling in relaying insulin action onto brain access of Ex-4_VT^750^, we used similar approaches as described in Figure 4. We found that, while VEGF injection (0.1 mg/kg) enhanced Ex-4_VT^750^ signal in the hypothalamus, administration of the VEGF-R antagonist Axitinib (25 mg/kg) prevented the action of insulin on Ex-4_VT^750^ access to this region (**Figure 5A**). Next, we generated mice lacking *Vegfa* in ME tanycytes (Langlet et al., 2013a). Vegfa^lox/lox^ mice were injected in the 3rd ventricle with either saline (Saline:;*Vegf^f/fx^*, control group) or a chimeric fusion protein Tat-Cre (Tat-Cre:;*Vegfa^f/f^*) to selectively ablate *Vegfa* in tanycytes (Tanycyte^ΔVegfa^ mice, **Figure 5B**). Control and Tanycyte^ΔVegfa^ mice received a 2-days treatment period consisting of daily injections (2:00 pm) of Ex-4 (120 nmol/kg, red), insulin (20 nmol/kg, blue), or a mix of insulin+Ex-4 (20 nmol/kg, 120 nmol/kg, green) (**Figure 5C**). A 3-days baseline period was first acquired for all groups (n=5 per group) and presented as reference (black, grey, **Figure 5B, C, D, E**). As previously observed, acute administration of Ex-4 and insulin oppositely impacted fatty acid oxidation (**Figure 5D, E**) in both control and Tanycyte^ΔVegfa^ mice. In control mice, the mix of insulin+Ex-4 induced a significant rebound in fatty acids oxidation able to oppose insulin-mediated decrease in fatty acids oxidation (**Figure 5D, F**), similarly to what we previously observed **(Figure 1E**). However, in Tanycyte^ΔVegfa^ mice, the metabolic consequences of Ex-4 and insulin co-administration was not different from that of insulin itself (**Figure 5E, G**). Importantly, the dose of insulin used in these experiments had a similar blood-glucose lowering action in both control and Tanycyte^ΔVegfa^ mice (data not shown). (**Figure S5);** however, when injected in combination with insulin, Ex-4 slightly opposed the increase in feeding in response to hypoglycaemia in control but not Tanycyte^ΔVegfa^ mice (**Figure S5).** Overall, these results suggest that tanycytic *Vegfa* relay, at least in part, the antagonistic action of insulin and Ex-4 onto fatty acids oxidation possibly by mediating local CVO’s adaptive response to hypoglycaemia and Ex-4 brain access.

**Figure 5.**
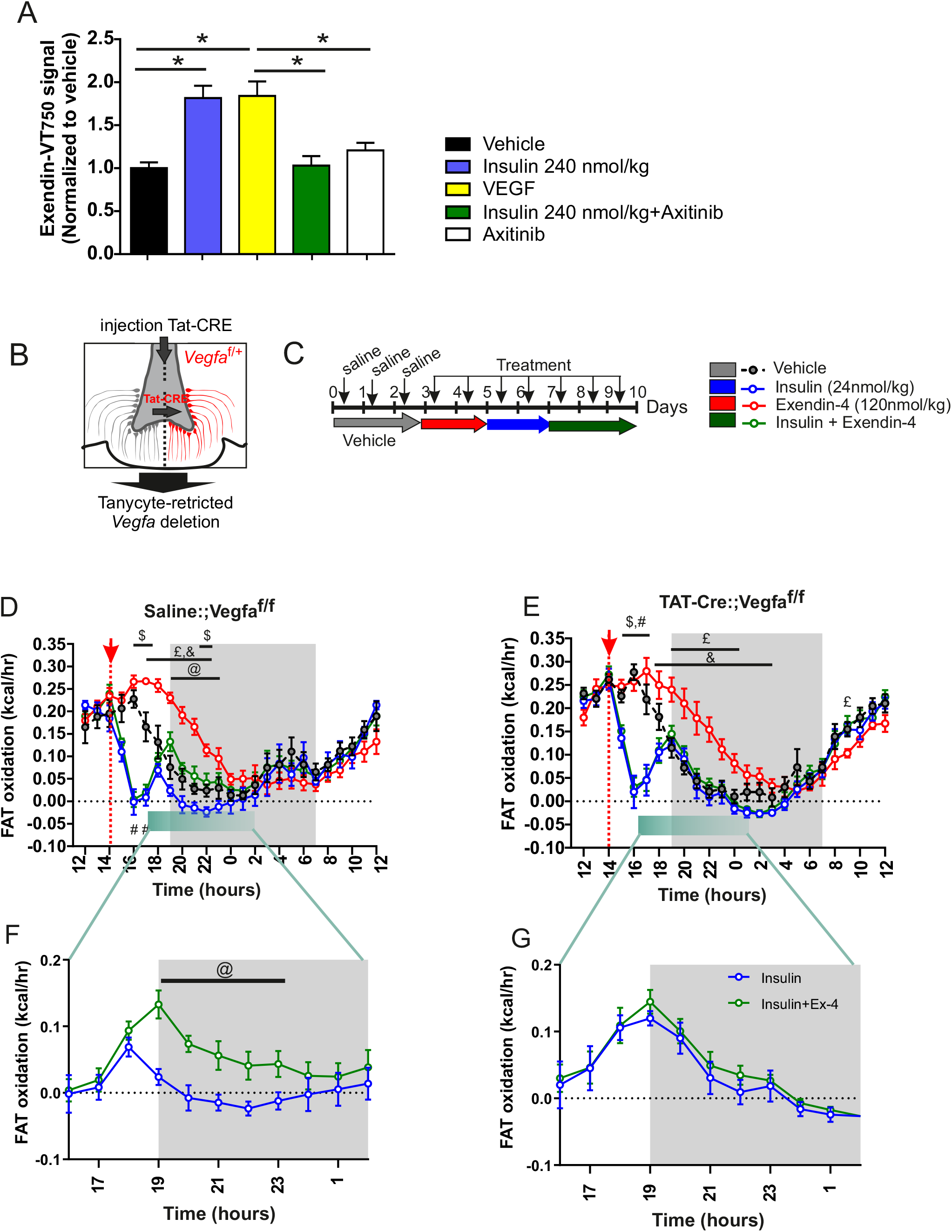
Metabolic action of Exendin-4 involves Tanycytes-born VEGF. (**A**) Quantification of hypothalamic Exendin-4_VT^750^ fluorescent signal normalized to vehicle in combination with IP injection of VEGF (0.1 mg/kg), Insulin (240 nmol/kg) +VEGF, VEGF receptor antagonist Axitinib (25 mg/kg) or Axitinib alone. n=5-17 per each group. **\***, indicated P<0.05, vehicle *vs* insulin. Data are expressed as mean ± SEM. **(B)** Model for tanycyte-restricted invalidation of *Vegfa* and (**C**) experimental set up for metabolic efficiency characterization of control (Vegfa^lox/lox^; 3rd ventricle injection of saline) and Tanycyte^ΔVegfa^ mice (Vegfa^lox/lox^; 3rd ventricle injection of TAT-CRE) in response to daily i.p. saline injection (baseline, grey) followed by a 2-days treatment period consisting of a daily injection (2:00 pm) of Exendin-4 (120 nmol/kg, red), followed by insulin (20 nmol/kg, blue), and 3 days of mix of insulin+Exendin-4 (20 nmol/kg, 120 nmol/kg, green). Graphs represent averaged values for all conditions (**D, E**) and close up of post-injection period for insulin (blue) and Insuline+Exendin-4 (green) average of a treatment days for (**F, G**). N=5 in each group. Data are expressed as mean ± SEM. $ indicates P value<0.05, insulin vs vehicle. £: indicates P value<0.05, Ex-4 vs vehicle. #: indicates P value<0.05, Insulin+Ex-4 vs vehicle. &: indicates P value<0.05, Insulin+Ex-4 vs Ex-4 @: indicates P value<0.05, Insulin+Ex-4 vs Insulin. Data are expressed as mean ± SEM. $ indicates P value<0.05

### The coupling between hypoglycaemia and brain access of Ex-4 is impaired following exposure to caloric-rich diet

We next explored the consequence of high-fat high-sugar (HFHS) exposure on the coupling between glucoprivation and brain access of Ex-4. In these conditions, adult male C57/Bl6J mice were exposed to a 5 or 10 weeks HFHS regimen and in each condition the ability of a single injection of insulin (240 nmol/kg) to promote BBB passage of Ex-4_VT^750^ was assessed. Mice exposed to HFHS during 5 and 10 weeks had a significant increase in body weight compared to chow-fed controls (**Figure 6A**). Insulin (240 nmol/kg) was able to trigger a similar decrease in blood glucose in both 5 and 10 weeks HFHS groups (**Figure 6B**). However, despite similar decrease in blood glucose, the ability of hypoglycemia to promote Ex-4_VT^750^ access to the brain was drastically decreased between 5 and 10 weeks HFHS exposure (**Figure 6C, D**) whereas in the hindbrain, after 5 weeks of HFHS diet, insulin-induce hypoglycaemia was still able to facilitate Ex-4 passage, action blocked in mice fed for 10 weeks (**Figure 6C, E**). The correlation between blood glucose changes in response to insulin and Ex-4_VT^750^ signal intensity revealed that in both hindbrain and hypothalamic structures the coupling between hypoglycaemia and brain access of GLP-1R agonists was impaired with 10 weeks of HFHS exposure (**Figure 6F, G**).

**Figure 6.**
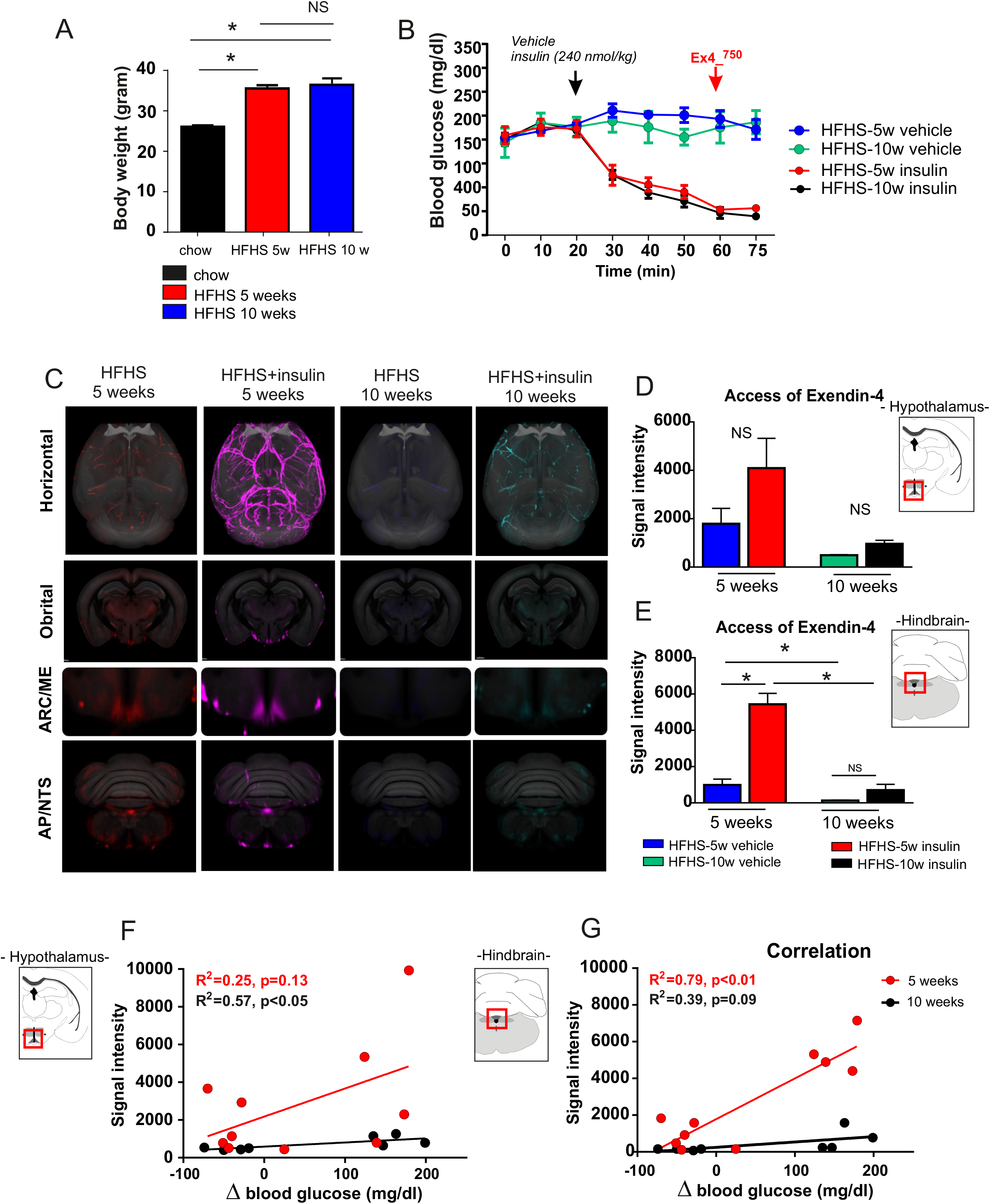
Exposure to energy-dense food leads to uncoupling between hypoglycemia and Exendin-4 brain access. (**A**) Body weight increase of mice exposed to 5 or 10 weeks of high fat high sucrose diet. (**B**) Change in blood glucose upon peripheral injection of insulin (240 nmol/kg) followed by the injection of fluorescent Exendin-4_VT^750^ (120 nmol/kg). (**C**) Horizontal, orbital planes digital section acquired with whole brains light sheet microscopy of fluorescent signals are overlaid onto the Common Coordinate Framework version 3 (CCFv3) template from the Allen Institute of Brain Science (Oh et al., 2014) Exendin-4_VT^750^ signal in the brain of mice after 5 or 10 weeks HFHS diet treated with vehicle or insulin (240 nmol/kg). (**D, E**) Quantification of Exendin-4_VT^750^ signal intensity in the hypothalamus (**D**) or hindbrain (**E**) structures. (**F, G**) Correlation of blood glucose levels (mg/dl) change upon insulin injection fluorescent signal intensity of Exendin-4_VT^750^ in (**F**) hypothalamic and (**G**) hindbrain region. n=5-6 minimum in each group. **\***, indicated P<0.05. Data are expressed as mean ± SEM.

### Blood glucose negatively correlates with central insulin content in humans

Taken together our results strongly suggest that, within a physiological range, acute perturbations of blood glucose promote rapid alterations in local brain access and action of Exendin-4. Indeed, we found that either systemic insulin-induced hypoglycaemia or central neuroglucopenia (2-DG injection) promoted accelerated Ex-4 entry in hypothalamic and hindbrain structures. We next explored the translatable aspect of this phenomenon in humans. We reasoned that this mechanism, in which lower plasma glucose leads to higher blood-to-brain passage of peptide hormones, could potentially also affect other circulating energy-related signals. In a cohort of non-obese individuals (N=140, characterization in Supplementary Table 2), we addressed the correlation between plasma glucose and the ratio of central CSF and systemic plasma insulin that reflects blood-to-brain insulin transport. When plasma glucose in fasted patients was regressed onto CSF/plasma insulin levels, we found a robust inverse correlation (p<0.001) showing that the lower the blood glucose the higher the ratio between central/systemic insulin (**Figure 7**). This result supports the notion that lower plasma glucose acts as a gate-keeper for central transport/access of blood-borne energy-related peptides also in humans. Importantly, while there was a negative correlation between the ratio of insulin in CSF/serum and body mass index (BMI) (p= 0.0007, r2=-0.079), the correlation between insulin in CSF/serum and glucose was independent of BMI and remained significant after adjustment for BMI (p<0.0001).

**Figure 7.**
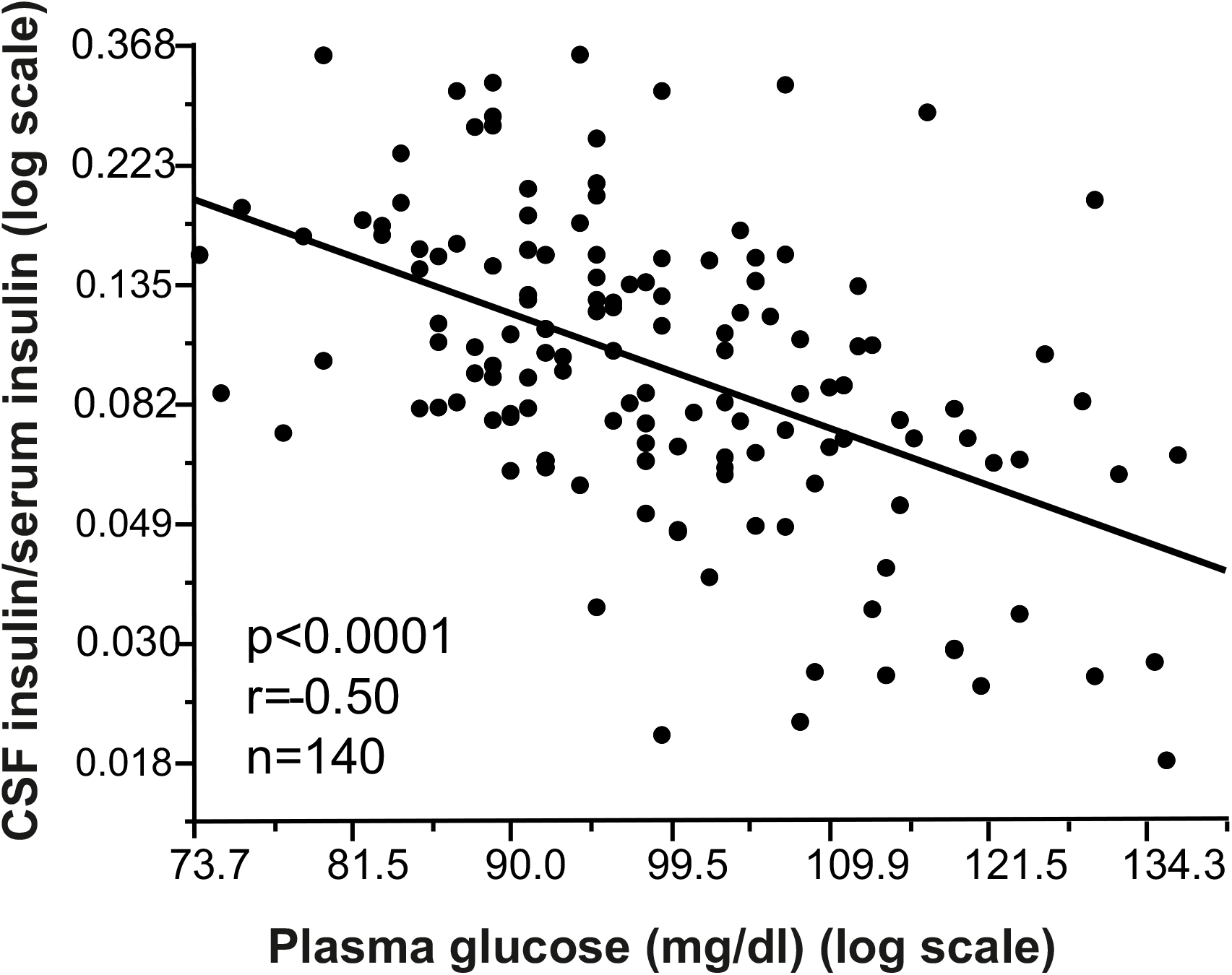
Central insulin content in human correlated with plasma glucose. Correlation of CSF/serum insulin ratio with plasma glucose in 140 human subjects. (P<0.0001, r = 0.50).

## DISCUSSION

The CNS has been identified as a primary site of action on energy expenditure and food intake for GLP-1R agonists (Baggio and Drucker, 2014). As both acute and chronic fluctuations in blood glucose have been shown to regulate adaptive changes in the BBB at CVO’s, these mechanisms potentially lead to modified brain access of circulating energy-related signals. We hypothesized that the centrally-mediated action of the GLP-1R agonists Exendin-4 on body-weight loss is influenced by either hypoglycemia and possibly through a direct central action of insulin.

We first showed that, when co-administered with insulin, Ex-4 opposes the action of insulin onto fuel partitioning towards glucose utilization by promoting a shift towards increased fatty acid oxidation (**Figure 1**). Using fluorescent labelled molecules, we were able to visualize that both insulin and Ex-4 can access ARC neurons and confirmed that the insulin-mediated signaling cascade (phospho-Akt) was rapidly initiated in the ME/ARC area (**Figure S2**). In an early time point after Ex-4 injection, we found an accumulation of Ex-4 signal in tanycytes residing in the ME but not in AP. This was associated with increased mRNA content for GLP-1R and InsR signaling in tanycytes of the ME/ARC compared to tanycytes in the AP/NTS region, suggesting that the InsR/GLP-1R tanycytic signaling is fostered in the ME/ARC (**Figure 2**).

Surprisingly, peripheral insulin injection accelerated the access of fluorescently labelled Exendin-4 in most CVOs including ME/ARC and AP/NTS regions (**Figure 3A and S3**).

In fact, we discovered that the degree of hypoglycemia, rather than insulin itself, correlated with an increased fluorescent Ex-4 signal in both the hypothalamus and the hindbrain (**Figure 3C-H**). This suggests that insulin signaling onto ME/ARC tanycytes did not confer a specific advantage for GLP-1R agonist access to the brain via CVOs but rather that other mechanisms might be involved.

The response of brain CVOs to hypoglycemia is, at least in part, relayed by central glucoprivic sensing, since 2-DG-induced glucopenia promoted rapid adaptive distribution of Ex-4 in hypothalamic structures (**Figure 4 G-I**) followed by change in hypothalamic and hindbrain Ex-4 abundance (**Figure 4 A-F**).

Using pharmacological and genetic approaches, we show that tanycytic VEGF-A release is an important mediator in the adaptive response of the BBB at CVOs to glycemic changes and the consequent action of Ex-4 on fatty acid oxidation (**Figure 5**).

Additionally, we demonstrated that exposure to an energy-rich diet induces a progressive defect in the adaptive access of Ex-4 to the brain in response to insulin-mediated glycemic changes (**Figure 6**). These results raise the possibility that obesity and/or high-fat feeding alters the ability of tanycytes to respond to acute changes during glycemia. Finally, we provide evidence that in humans, glycaemia also correlates with blood-to-brain entry of the energy-related signals: insulin.

Altogether, these results support the notion that acute changes in plasma glucose, detected at the central level, induce a rapid adaptive modification of brain access and action of energy-related signals. In the framework of insulin/GLP-1R agonist co-treatment, our data supports a model where tanycytic VEGF-A relay the action of insulin-mediated hypoglycemia onto ME/ARC entry and action of GLP-1R agonist. Indeed, to mediate their action, circulating signals need to reach neurons that lie behind the BBB. First order neurons, such as AgRP- and POMC-neurons, have an exquisite location close to the capillary loops arising from the ME (Schaeffer et al., 2013), with a tightly regulated permeability that allows central access of circulating molecules via CVOs (Banks, 2019; Prevot et al., 2018). Permeability of the ME vascular loops reaching the ARC is tightly regulated by tanycytes, which are glucose-sensing cells (Frayling et al., 2011) and one of the main sources of VEGF-A in the ME (Jiang et al., 2020; Langlet et al., 2013a; Langlet et al., 2013b). The tanycytes regulate the direct access of blood-borne molecules to ARC neurons by controlling capillary fenestration. Hence, the microenvironment determined by the structure of the local capillaries and tanycytes-mediated signaling might confer to each CVO an idiosyncratic and distinct ability to integrate changes in nutrients availability and adapt blood-to-brain passage of metabolic signals. This hypothesis is in line with a recent study by Fortin and colleagues which demonstrates that AP is not required for food intake and body weight-reducing effects of acutely delivered liraglutide, while GLP-1R signaling onto NTS neurons mediates part of the acute anorectic effects of liraglutide (Fortin et al., 2020).

In our study, we observed a region-specific differential expression of GLP-1R/InsR between hypothalamic and hindbrain tanycytes (**Figure 2**), highlighting a potentially more prominent role for hypothalamic tanycytes in the potential access of GLP-1R agonists in the ARC, while adaptive responses to fenestrated capillaries could be responsible for changes in hindbrain accessibility. In support of this view, we observed that hypothalamic tanycytic VEGF-A signaling is required for the insulin-promoted Ex-4 access and distribution in the ARC and for the GLP-1R agonist action on peripheral lipid-substrate utilization (**Figure 5D-G**).

In that sense, our work suggests that hypothalamic access is important for long-term fat mobilization induced by GLP-1R agonists, in good agreement with the work of Secher et al who identified the ARC as an important site for the long-term, weight-reducing action of liraglutide (Secher et al., 2014). However, our results do not preclude the possibility that GLP-1R agonists also act on other neural substrate in the NTS (Fortin et al., 2020). Indeed, our study together suggest that glycemic change can modulate the passage of GLP-1R agonist in hindbrain and hypothalamic structure through tanycyte-dependent and independent mechanisms, and throughout different timeframe.

In that view, GLP-1-induced anorectic responses could be mediated by hindbrain structures while change in peripheral glucose would promote adaptive change in ME, with a consequence on whole body fat oxidation through local action onto hypothalamic neural network.

The action of GLP-1R on both neural substrates ultimately acting in synergy to mediate the body weight loss action of GLP-1R agonist throughout different time scale. The contribution of arcuate neurons to the action of GLP-1R agonists on lipid substrate utilization is well in line with previous studies showing that, 1) peripheral and intra-arcuate injection of Ex-4 can oppose the decrease in lipid oxidation induced by centrally delivered ghrelin (Abtahi et al., 2016) and, 2) GLP-1R agonists enhance triglyceride clearance in human (Whyte et al., 2018).

Overall, these observations fit with a concept where appetite-suppression and fat mobilization resulting from a combinatorial action of peripherally administered GLP-1R agonists and insulin both contribute to weight loss. It is probable that the timing by which GLP-1R agonists reach hypothalamic and brainstem targets will be influenced by adaptive mechanisms governing peptide entry into these regions through respective distinct CVOs. One can therefore envision an orchestrated response pattern with a first acute response evoked in the vagal nerve and in the AP/NTS, followed by second, more sustained, increase in energy expenditure/favored lipid utilization through action on ARC neurons. Our data also highlight that the increase in fat oxidation induced by GLP-1R agonist can oppose the lipogenic action of of insulin. In theory, in a context of both excessive body weight and uncontrolled glycemia, achieving glucose control while limiting insulin-mediated lipogenesis and body weight gain would probably have beneficial output on metabolic syndrome (Muller et al., 2019).

The present study suggests that glucose fluctuations are instrumental in adaptive changes of peptide entry through CVOs and consequent central actions of GLP-1R analogs. In that regard, one would expect that the timing of insulin and GLP-1R agonist co-treatment might be important to harness the full benefit of this pharmacological approach. In condition of energy plethora and obesity or, at the opposite, after repeated hypoglycemia and partial loss of glucose awareness, it is possible that this adaptive response of the peptide entry into the brain to glucose change is blunted. Indeed, we found that exposure to HFHS diet decreases the CVO adaptive response to hypoglycemia leading to blunted Ex-4 entry in the brain.

This mechanism might also be important for endogenous peptide signals to reach the target neuronal cell population, especially for metabolic signals characterized by short plasma half-life such as GLP-1 and insulin. Langlet and colleagues first described that plastic adaptations of the ME/ARC to glucoprivation involves VEGF-A (Langlet et al., 2013a). Our study is in accordance with these observations and provides further evidence for the implication of tanycytic-VEGF-A in local BBB remodeling. However, it is also possible that systemic VEGF-A plays an important role. Indeed, insulin-mediated hypoglycemia increased serum VEGF-A in humans (Dantz et al., 2002), whilst no modifications were observed under hyperinsulinemic-euglycemic clamp (Loebig et al., 2010). This observation suggests that, similar to what we observed in our study (**Figure 3**), glucose dynamics rather than insulin itself may be responsible for plasma VEGF-A alterations. In line with this hypothesis, studies using euglycemic-clamp revealed a relationship between elevated plasma VEGF-A in both obese humans (Loebig et al., 2010) and after exposure to high-fat diet (Schuler et al., 2018), independent of insulin sensitivity. These striking similarities in the responsiveness of both systemic and tanycytic VEGF-A suggest that conserved nutrient-sensing mechanisms, yet to be identified, might govern VEGF-A synthesis in tanycytes and peripheral organs. In conditions of nutrient overload or uncontrolled plasma glucose, systemic VEGF-A could bypass tanycytic-born VEGF-A signaling, thus leading to local adaptations in the ARC. A recent study shows that activation of melanin-concentrating hormone (MCH) neurons in the lateral hypothalamic area also regulates ME/ARC vessels’ permeability possibly via the release of VEGF-A (Jiang et al., 2020). However, because insulin has been shown to have a strong inhibitory effect on MCH neurons (Silva et al., 2009), they are unlikely to be involved in the hypoglycemia/insulin-promoted facilitation of the access of GLP1-R agonists to the ARC.

A limitation of our study is that we do not provide a mechanism by which HFHS exposure uncouples brain response to hypoglycemia. Previous reports have argued that exposure to energy-dense food and altered nutrient sensing can impair tanycyte function (Balland et al., 2014; Langlet et al., 2013a) and also trigger inflammatory-like response in the hypothalamic region in humans and rodents (Andre et al., 2017; Arruda et al., 2011; Thaler et al., 2012) (Banks, 2003; Banks et al., 2004; Banks et al., 1999). Most of these mechanisms may play an important role in maladaptation of BBB responsiveness and the consequent dysregulation of brain-entry and action of circulating signals.

Overall, our study suggests that acute changes in plasma glucose within a physiological range, gates the level of access of energy-related peptide signals to specific neuronal populations that in turn serve as central controllers of body weight homeostasis by regulating appetite and substrate utilization. In support to our working hypothesis, we found that in non-obese humans, plasma glucose was negatively correlated with the ratio between circulating and central insulin (**Figure 7**), indicating an increase of central insulin entry when plasma glucose is low. Beyond the translational perspective, this observation in humans, together with the results obtained in rodents, suggest that achieving glycemic control is of crucial importance as it might directly alter brain access and response to important metabolic factors. Recent studies have highlighted defects in insulin entry into the brain in insulin-resistant humans (Heni et al., 2014) (Heni et al., 2015), promoting the concept that defective brain action of insulin might accelerate the progression of metabolic disease. In line with these observations, the work presented here reveals that uncontrolled glycaemia, could be a starting point in defective access for metabolic peptide signals to relevant neural populations regulating feeding behavior and energy expenditure. It is likely that such a situation may aggravate metabolic disease by fueling a vicious circle where increased appetite and decreased lipid utilization further perpetuates weight gain. While our study holds interesting avenue for better understanding and optimization of the action of circulating signals onto central regulation of metabolism, further studies are warrant to fully understand the mechanisms by which peptide access into the brain is modulated by glycemia change and how to circumvent/correct the default acquired through nutrient overload and obesity.

## EXPERIMENTAL PROCEDURES

### ANIMALS

All animal experiments were conducted in accordance with approved national regulations of both France and Denmark, which are fully compliant with internationally excepted principles for the care and use of laboratory animals, and with animal experimental licenses granted by either the Animal Care Committee University Paris Diderot-Paris 7 (CEB-14-2016) or the Danish Ministry of Justice. C57Bl6J male mice (10-18 weeks) were housed 5 per cage or individually when required for the experiment, in standard, temperature-controlled condition with a 12-hour-light/dark cycle. The mice had ad libitum access to water and regular chow, unless otherwise stated. tdTomato^loxP/+^ reporter mice were purchased from the Jackson Laboratories (Bar Harbor, ME) and Vegfa^loxP/loxP^ mice (Gerber et al., 1999) were a gift from N. Ferrara (Novartis).

#### Hyperinsulimic high/low glycemic clamp

Catheter was chronically inserted in the jugular vein under anesthesia and mice recovered for one week before starting the experiment. Mice were food deprived for 5 hours to establish stable blood glucose levels. Mice were connected through the jugular vein catheter in the clamp system and after 30 min injected with insulin bolus (100 mU/kg) and continuous maintained with 3.3 mU/kg/min Insulin (Novorupid, Flex pen, 8-9670-79-202-2), 2 ul/min speed with pump. Mice were either maintained on glucose levels between 140-160 mg/dl (high) or maintain on glucose levels between 70-80 mg/dl (low) controlled by 30% glucose solution connected to the clamp. For 140-160 mg/dl (high glucose), the pump was started 10 minutes after insulin bolus with speed at 4ul/min. For 70-80 mg/dl (low glucose), glucose perfusion only started when BG levels dropped to 60mg/dl within 30 minutes. Blood glucose was measured every 10 minutes during the entire experiment by tail cut and measured by Glucofix glucose analyzer. After approximately 90 min, when stable high or low glucose was established, mice were injected with fluorescent labelled Exendin4 (Exendin4_VT^750^, 120 nmol/kg, Novo Nordisk) 15 minutes prior to sacrifice, followed by fixation perfusion (10 ml Saline with 20 U/ml heparin and 10ml 10% Neutral Buffered Formalin) and dissection of the brain. Tissues were stored in 10% NBF 4°C until further processing.

#### 2DG-induced neuroglucopenia

Central neuroglucopenia was induced by 250 mg/kg of 2-deoxyglucose (Sigma) injected ip after 5 hour of food deprivation. ~10 min prior to sacrifice and 30 min after 2-DG injection, animals were injected with fluorescent Exendin-4 (Exendin4_VT^750^ or Exendin4_^Cy3^, 120nMol/kg) at early (15min) or late (40min) time points.

#### Compound and doses

Insulin VT^750^ was used i.v. at 240 nM/kg, and mice were sacrificed 10-15 sec or 15 min after injection. S961 (NOVO NORDISK) was used i.v. at 1060 umol/kg and injected 30 min prior insulin administration.

Recombinant mouse VEGF (RDS) was delivered i.p. at 60 ug/kg as previously described (Langlet et al., 2013a). Axitinib (in DMSO, LC Laboratories, France) was delivered i.p. at 25 mg/kg as described in (Langlet et al., 2013a). Exendin-4 VT^750^, Exendin-4 ^CY3^, Insulin VT^750^, liraglutide and insulin were kindly provided by Novo Nordisk.

#### Saline or Tat-CRE mediated central delivery

Procedure was identical to that described in (Langlet et al., 2013a; Peitz et al., 2002). Saline or a Tat-Cre fusion protein was stereotaxically (Peitz et al., 2002) infused into the 3rd ventricle (1.5 ml over 5 min at 2.1 mg/ml; anteroposterior, −1.7 mm; midline, 0 mm; dorsoventral, –5.6 mm) of isoflurane-anesthetized Vegfa^loxP/loxP^ mice to produce control or Tanycyte^ΔVegfa^ mice respectively. Tat-Cre fusion protein was injected in the 3rd or in the 4th ventricle (1.5 ml over 5 min at 2.1 mg/ml; anteroposterior, −6 mm; midline, 0 mm; dorsoventral, –4 mm) of tdTomato^loxP/+^ reporter mice.

#### Tanycyte isolation and PCR analysis

Fluorescence-Activated Cell-Sorting (FACS) and Real-Time PCR Analyses of tdTomato-positive cells were performed on cells sorted and collected from ME and AP explants microdissected from tdTomato^loxP/+^ mice following 3rd or 4th ventricle Tat-Cre injection and processed for quantitative RT-PCR as described in (Langlet et al., 2013a). List of primers used is shown as Supplementary table 1.

### IMAGING TECHNIQUES

#### Tissue processing for laser sheet microscopy analysis

For light sheet fluorescent microscopy, the tissues were dehydrated in increasing concentrations (50 vol%, 80 vol%, 96 vol%, 2 × 100 vol%) of tetrahydrofuran (Sigma), at 3 to 12 hours per step. The dehydrated sections were then cleared until transparent by incubating with 3 × dibenzylether (Sigma) for 1 to 2 days in total. For immunohistochemistry, tissues were immediately saturated in 20% sucrose solution in 4% PFA overnight at 4°C, followed by embedding and freezing in tissue teck. Brain transparisation was performed as previously described (Renier et al., 2014; Salinas et al., 2018).

#### Light sheet microscopy

Visualization of fluorescence was performed with a light sheet ultramicroscope coupled to a SuperK EXTREME (EXR-15) laser system (LaVision). The samples were scanned in 5-μm steps at excitation/emission settings of 620/700 nm for autofluorescence signal and 710/775 nm for specific signal of exendin4^750^ or insulin^750^. All samples were scanned with identical settings. Images were generated using Imaris Bitplane software (Imaris x64 7.5.1). The fluorescent signals shown in Figure 3 and Figure 6 are unmixed signals. The estimated auto-fluorescence contribution in the specific channel was calculated and removed based on ratios of voxel intensities between selected voxels in the autofluorescence recording and the corresponding voxels in the specific recording. The unmixing algorithm was written in Matlab (Release 2012b, MathWorks, Natick, Massachusetts, United States) and applied as a XT-plugin in Imaris (Release 7.6.5, Bitplane, Zurich, Switzerland).

#### Immunohistochemistry

Brains were sectioned into 30-μm cryosections covering the hypothalamus. All sections were collected and consecutively sampled on cryoslides and stored in −80C. Sections were blocked in TBS containing donkey serum or PBS/0.1% Triton X-100 and stained for Vimentin (BD pharmingen #550563, 1:2000) and phospo-Akt (CST #4060 1:50) in PBS/0.1% Triton X-100/0.2% BSA at 4°C overnight. Brain sections were counterstained with Hoechst nuclear stain (1:10,000 μl in dH_2_O). Images were obtained using a Leica TCS SP8 confocal microscope. Tissue processing: tissues are just after dissection saturated in 20% sucrose solution in 4%PFA overnight at 4°C, followed by embedding and freezing in tissue teck. GLP-1R antibody was produced by Novo Nordisk (REF)

#### Confocal microscopy

For hypothalamic Exendin-4 Cy3 signal quantification, mice were rapidly anaesthetized with gaz isoflurane and transcardially perfused with 4% (weight/vol.) paraformaldehyde in 0.1 M sodium phosphate buffer (pH 7.5). Brains were post-fixed overnight in the same solution and stored at 4°C. 30 μm-thick sections were cut with a vibratome (Leica VT1000S, France), stored at −20 °C in a solution containing 30% ethylene glycol, 30% glycerol and 0.1 M sodium phosphate buffer. Sections were processed as follows: Day 1: free-floating sections were rinsed in Tris-buffered saline (TBS; 0.25 M Tris and 0.5 M NaCl, pH 7.5).

Acquisitions for Exendin-4 fluorescent signal were performed with a confocal microscope (Zeiss LSM 510) with a color digital camera and AxioVision 3.0 imaging software. Images used for quantification were all single confocal sections. Photomicrographs were obtained with the following band-pass and long-pass filter settings: Cy3 (band pass filter: 560–615). The objectives and the pinhole setting (1 airy unit, au) remained unchanged during the acquisition of a series for all images. Quantification of fluorescent signal was performed using the ImageJ software taking as standard reference a fixed threshold of fluorescence. Insulin VT^750^

#### Total Protein Extraction and western blot Analysis

Mice were injected i.p. with insulin (60 nmol/kg, Novo Nordisk) or vehicle prior to brains were dissected by decapitation. Hypothalamic and MBH region were detected and isolated under microscope and were snap frozen immediate in liquid nitrogen. Proteins were extracted with tris-based lysis buffer (25 mM Tris pH7.4, 50 mM beta-glycerophosphate, 1% triton x100, 1.5 mM EGTA, 0.5 mM EDTA, 1mM sodium pyrophosphate, 1 mM sodium orthovanadate, 10ug/ml leucepeptin + pepstatin A, 10 ug/ml aprotin, 100ug/ml PMSI). Homogenates were incubated on ice for 30 min and then cleared by a 10 min centrifugation at 21,000 g. 20 ug of protein extracts were fractionated by SDS-PAGE and transferred to nitrocellulose membrane for western blot analysis. Primary antibodies were used at the following dilutions: 1:100 (phospho-Ak (ser473): CST #4060S), 1:500 (total-Akt: CST #9272), 1:1000 (phospho-ERK (T42/44), CST #9101), 1:5000 (beta actin: Sigma A1978).

### METABOLIC MEASUREMENTS

#### Acute insulin/Exendin-4 injection

Metabolic output and feeding were monitored on C57Bl6j male mice (10 weeks old) using indirect calorimetry (n=8 per groups) for a 2-days baseline acclimation consisting of a single saline injection (i.p at 2:00 pm), followed by a 4-days treatment period in which animals received an i.p injection of insulin (24 nmol/kg), Exendin-4 (120 nmol/kg) or a combination of both (Ex-4 120 nmol/kg+Insulin 24 nMol/kg). After completion of the analysis, animal received an acute injection of fluorescent labelled Ex-4 (Exendin4_VT^750^, 120nMol/kg, Novo Nordisk) 15 minutes prior to sacrifice, followed by fixation perfusion (10 ml saline with 20U/ml heparin and 10ml 10% Neutral Buffered Formalin) and dissection of the brain. Tissue were stored in 10% NBF 4°C until further processing. Tanycyte^ΔVegfa^ and control mice were subjected to a 3-day saline baseline followed by consecutive, 2-days regimen of Exendin-4, 2-days of and insulin and 3-days Insulin +Exendin-4 injection. AgRP-ablated and control naïve mice were monitored in the same paradigm for a 3-days saline (i.p) injection period followed by a 7-days regimen consisting of daily injection of liraglutide (0.1mg/kg, s.c, at 14h00).

#### Metabolic efficiency analysis

Mice were monitored for whole energy expenditure (EE) or Heat (H), oxygen consumption and carbon dioxide production, respiratory exchange rate (RER = VCO2 / VO2, where V is a volume), and locomotor activity using calorimetric cages with bedding, food and water (Labmaster, TSE Systems GmbH, Bad Homburg, Germany).

Ratio of gases is determined through an indirect open circuit calorimeter (for review (Arch et al., 2006; Even and Nadkarni, 2012)). This system monitors O2 and CO2 concentration by volume at the inlet ports of a tide cage through which a known flow of air is being ventilated (0.4 L/min) and compared regularly to a reference empty cage. For optimum analysis, the flow rate is adjusted according to the animal weights to set the differential in the composition of the expired gases between 0.4 and 0.9% (Labmaster, TSE Systems GmbH, Bad Homburg, Germany).

The flow is previously calibrated with O2 and CO2 mixture of known concentrations (Air Liquide, S.A. France). Oxygen consumption, carbon dioxide production and energy expenditure were recorded every 15 min for each animal during the entire experiment. Whole energy expenditure was calculated using the Weir equation respiratory gas exchange measurements (3). Fatty acid oxidation was calculated from the following equation: fat ox (kcal/h) = energy expenditure X (1-RER/0,3) according to Bruss et al (Bruss et al., 2010).

Food and water consumption were recorded as the instrument combined a set of highly sensitive feeding and drinking sensors for automated online measurement. Mice had free access to food and water *ad libitum*. To allow measurement of every ambulatory movement, each cage was embedded in a frame with an infrared light beam-based activity monitoring system with online measurement at 100 HZ. The sensors for gases and detection of movement operate efficiently in both light and dark phases, allowing continuous recording.

Estimation of resting metabolism according Péterfi et al. (Peterfi et al., 2018) was made on the following basis: data points of EE were considered to be the best estimation of resting metabolism when spontaneous activity during the previous 30 minutes was less than 1-5% of the highest daily value and that animals ate less than 0,1g of food within the last hour.

Mice were monitored for body weight and composition at the entry and the exit of the experiment. Body mass composition (lean tissue mass, fat mass, free water and total water content) was analyzed using an Echo Medical systems’ EchoMRI (Whole Body Composition Analyzers, EchoMRI, Houston, USA), according to manufacturer’s instructions. Briefly, un-anesthetized mice were weighed before they were put in a mouse holder and inserted in MR analyzer. Readings of body composition were given within 1 min.

Data analysis was performed on excel XP using extracted raw value of VO2 consumed, VCO2 production (express in ml/h), and energy expenditure (Kcal/h). Subsequently, each value was expressed either by total body weight or whole lean tissue mass extracted from the EchoMRI analysis.

#### Analytical procedures in humans

Cerebrospinal fluid (CSF) and blood (serum and NaF-plasma for glucose measurements) collection was performed under fasting conditions between 8:00 and 10:00 a.m. and subjects with normal CSF cell counts on the XN-10 hematology analyzer (Sysmex, Norderstedt, Germany) were included in the study. We included patients with Parkinson’s disease (PD) as this enabled a highly structured, standardized and fast collection of specimen in the frame of the prospective ABCPD study (Sulzer et al., 2018). Such a high quality collection especially of CSF is almost impossible in comparably large cohorts of healthy subjects, and we are not aware of any literature reporting about glucose pathway alterations in PD. asma and CSF glucose were measured immediately after collection on the ADVIA XPT clinical chemistry analyzer (hexokinase method). Plasma and CSF insulin levels were determined by an ADVIA Centaur XPT chemiluminescent immunoassay system after testing linearity and recovery for insulin in CSF (both instruments from Siemens Healthineers, Eschborn, Germany). The variation of this assay in CSF at an insulin concentration of 8 pmol/l was 12 %. The ethics committee of the Medical Faculty of the University of Tuebingen approved the study (686/2013BO1), and the study was performed in accordance with the Declaration of Helsinki.

### STATISTICS

All data were entered into Excel 5.0 or 2003 spread sheets and subsequently subjected to statistical analyses using GraphPad Prism or Statview Software. Statistical significance was set to *P* < 0.05. Statistical evaluation of the data was carried out using 1- or 2-way repeated-measures ANOVA, with Fisher’s post-hoc analysis (1-way) or Bonferroni post-hoc analysis (2-way) and student t-test between control and treatment groups in cases in which statistical significance was established.

## Supporting information

Supplemental information includes 5 figures, 2 tables and supplemental experimental procedures

## SUPPLEMENTAL INFORMATION

Supplemental information includes 5 figures, 2 tables and supplemental experimental procedures

## ACKNOWLEDGMENTS

This work was supported by a collaborative research grant from NOVO NORDISK and the Université de Paris. We acknowledge funding supports from the Centre National la Recherche Scientifique (CNRS), The Université de Paris. We thank Olja Kacanski for administrative support, Isabelle Le Parco, Ludovic Maingault, Angélique Dauvin, Aurélie Djemat, Florianne Michel, Magguy Boa and Daniel Quintas for animals’ care and Sabria Allithi for genotyping. We acknowledge the technical platform Functional and Physiological Exploration platform (FPE) of the Université de Paris, BFA, UMR 8251, CNRS, Paris, France and the animal core facility “Buffon” of the Université de Paris/Institut Jacques Monod. DHMC and RH received support from the National Research Agency ANR-15-CE14-0030-01:” Nutritpathos” and ANR-17-CE37-0007-02 “METACOGNITION” respectively.

---

**Figure S1.**
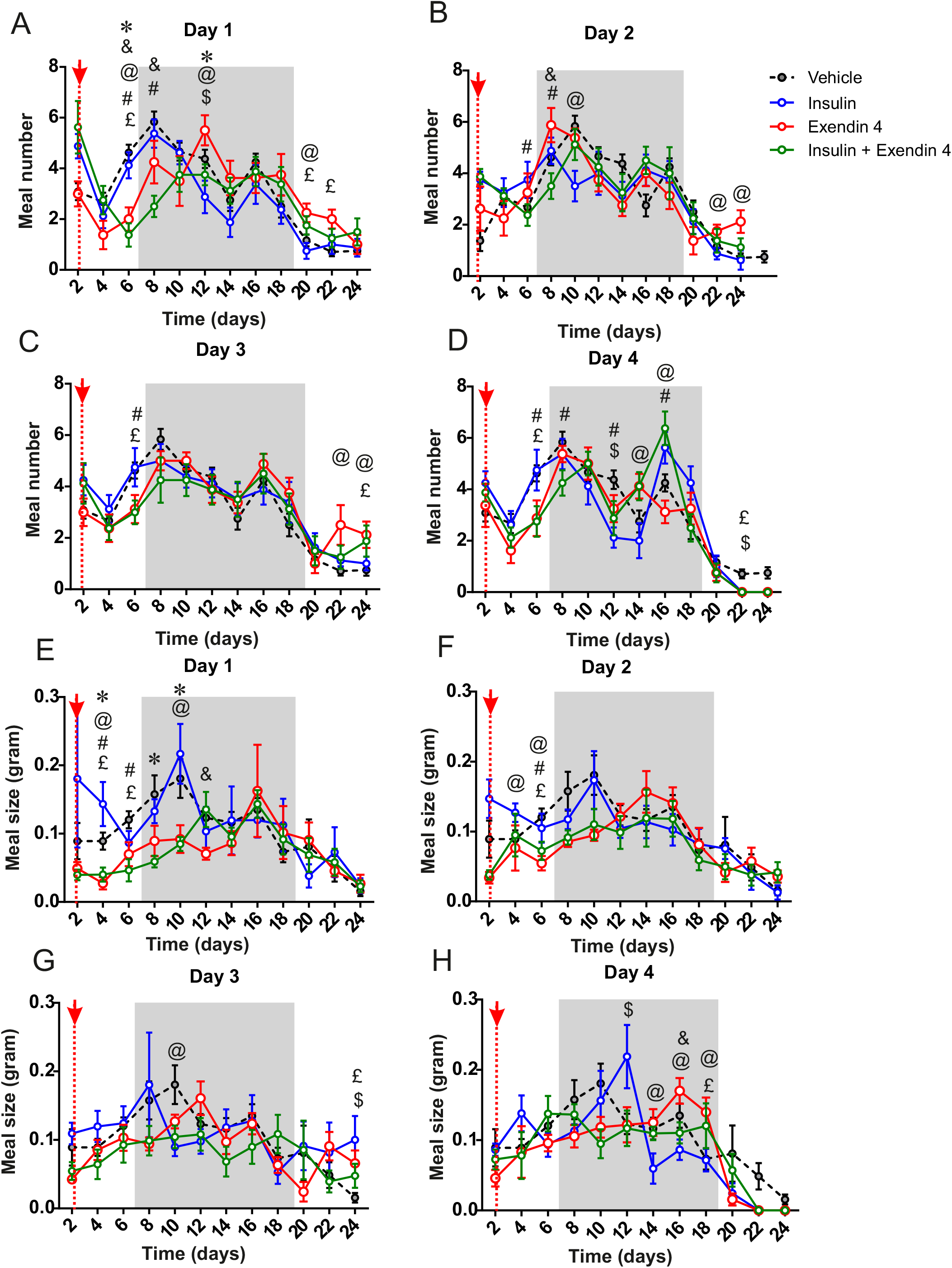
Meal ultrastructure analysis during insulin and Exendin-4 co injection. Mice received a daily IP saline injection (baseline, grey) followed by a 4-days treatment period consisting of a daily injection (14:00 pm) of insulin (20nmole/kg, blue), Exendin-4 (120nmole/kg, red) or mix of insulin (20nmole/kg) + Exendin-4 (20nmole/kg, 120nmole/kg, green). Graphs represent an average of a 3-days baseline (grey) or treatment periods 3-days treatment for meal number (**A-D**) or meal size (**E-H**). N=8 in each group. Data are expressed as mean ± SEM. **\*** indicated P value<0.05. $ indicated P value<0.05, ins vs vehicle. £: indicated P value<0.05, Ex-4 vs vehicle. #: indicated P value<0.05, Insulin+Ex-4 vs vehicle. &: indicated P value<0.05, Insulin+Ex-4 vs Ex-4. @: indicated P value<0.05, Insulin vs Ex-4 .*: indicated P value<0.05, Insulin+Ex-4 vs Ex-4

**Figure S2.**
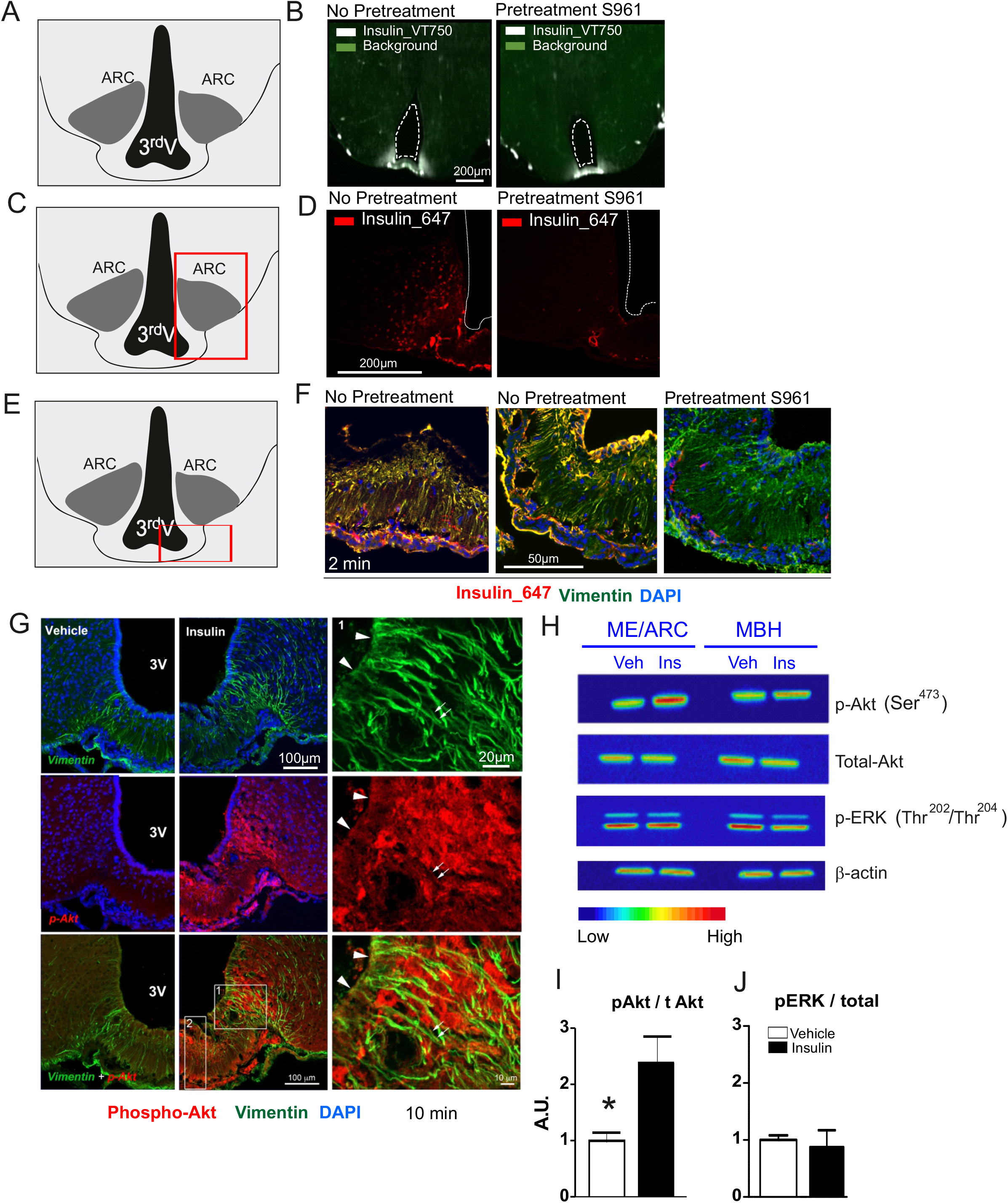
Peripheral Insulin access and action on ME/ARC hypothalamic regions. (**A-F**) Representative photomicrographs of median eminence (ME) and arcuate (ARC) hypothalamic regions acquired with light sheet microscopy after peripheral injection of fluorescent insulin (**B**, Insulin-VT^750^, **D-F** Insulin_Alexa^647^, 240 nmol/kg) with or without pre-treatment with the insulin receptor antagonist S961. (**F**) Fluorescent signal for i.v. injected Insulin_Alexa^647^ (240 nmol/kg) (red) was associated with immunochemical detection of vimentin (green) and DAPI (blue). (**G**) Representative immunohistochemical analysis of phospho-Akt (red), vimentin (green) and DAPI (blue) in ME tanycyte projections. (**H, I**) Representative western blot analysis (**H**) in heatmap colors and signal quantification (**I, J**) of phospho-Akt and phospho-ERK in mediobasal hypothalamus (MBH) and ME/ARC punches 10 min after i.v. insulin administration (iv 60 nmol/kg). n=3 in each group. **\***, indicates P<0.05, vehicle *vs* insulin. Data are expressed as mean ± SEM.

**Figure S3.**
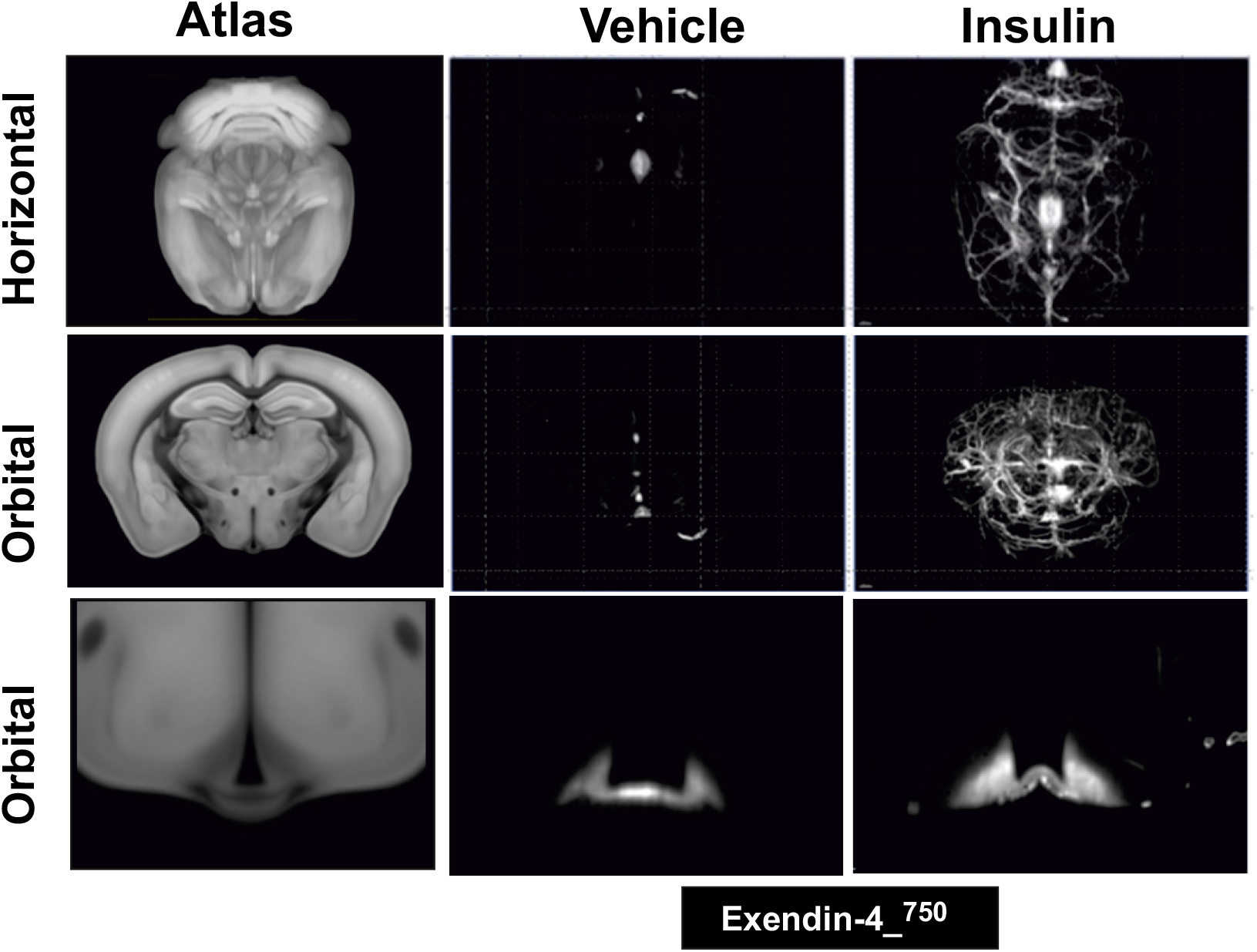
Insulin potentiates the access of GLP-1 analogue Exendin-4 into the brain. Selection 2D-planes whole brain light sheet scanning to visualize fluorescent signal of peripherally injected Exendin-4_VT^750^ (120 nmol/kg) together with intraperitoneal (ip) injection of vehicle or insulin (240 nmol/kg). The picture represents horizontal and coronal planes of the brain with magnification of ME/ARC structures compared to Allen’s brain atlas (Lein et al., 2007).

**Figure S4.**
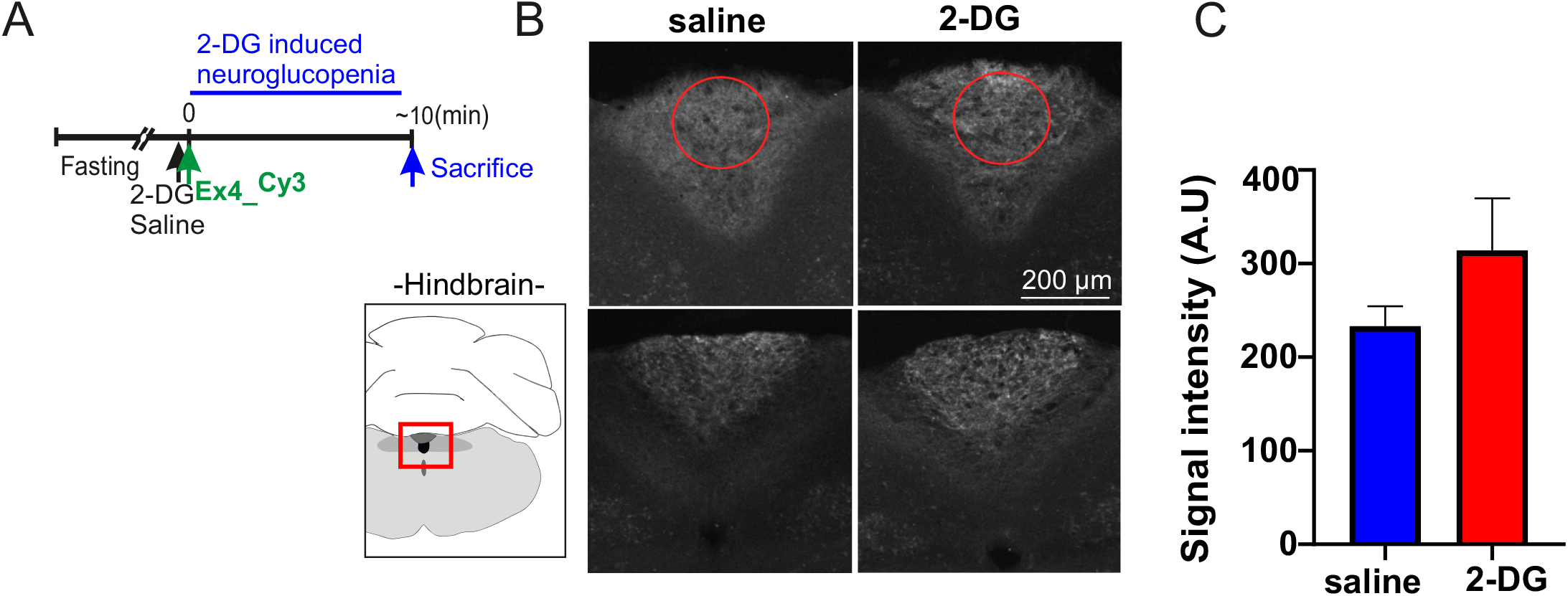
Central neuroglucopenia increases Exendin-4 access to the brain. (**A**) Experimental set up for concomitant injections of 2-DG (250 mg/kg) and Exendin-^Cy3^ (120 nMol/kg) ~10 min prior to sacrifice. (**B**) Representative photomicrograph for Exendin-^Cy3^ fluorescent distribution and (**C**) signal quantification in the middle of the AP (red circles) of the AP ~15 min after 2-DG bolus injection. n=5-9 minimum in each group. **\***, indicated P<0.05. Data are expressed as mean ± SEM.

**Figure S5.**
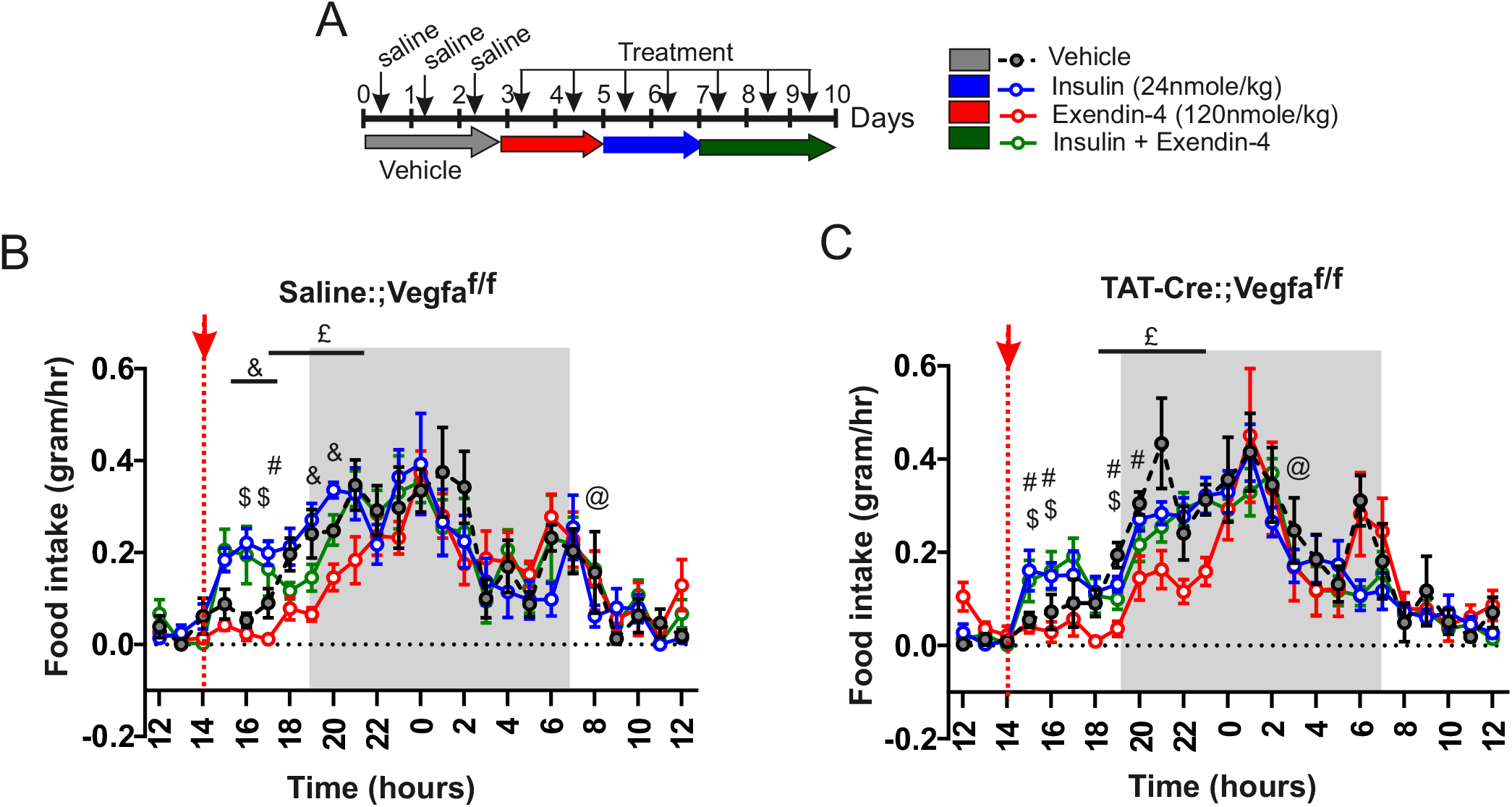
Role of Tanycyte-born VEGF in glucose-mediated brain entry and action of Exendin-4. (**A**) Experimental set up for metabolic efficiency characterization of control (Vegfa^lox/lox^; 3rd ventricle injection of saline) and Tanycyte^ΔVegfa^ mice (Vegfa^lox/lox^; 3rd ventricle injection of TAT-CRE) receiving a daily IP saline injection (baseline, grey) followed by a 2-days treatment period consisting of a daily injection (14:00 pm) of Exendin-4 (120nmole/kg, red), followed by insulin (20nmole/kg, blue), and 3 days of mix of insulin (20nmole/kg) + Exendin-4 (20nmole/kg, 120nmole/kg, green). (**B, C**) Averaged food intake over the treatment period in control (Vegfa^lox/lox^, with 3rd ventricle injection of saline, **B**) or Vegfa^lox/lox^, 3rd ventricle injection of TAT-CRE, **C**) or . N=5 in each group. Data are expressed as mean ± SEM. $ indicated P value<0.05, insulin vs vehicle. £: indicated P value<0.05, Ex-4 vs vehicle. #: indicated P value<0.05, Insulin+Ex-4 vs vehicle. &: indicated P value<0.05, Insulin+Ex-4 vs Ex-4 @: indicated P value<0.05, Insulin+Ex-4 vs Insulin.

