## Supplemental information includes 5 figures, 2 tables and supplemental experimental procedures for "Acute changes in systemic glycaemia gate access and action of GLP-1R agonist on brain structures controlling energy homeostasis"

**Supplementary material**

**Methods**

**Meal ultrastructure**

Meals Ultra structure (meal number, size and frequency, bouts and inter meal intervals) were defined as described in as define by Gaetani et al. (2003) (Gaetani et al., 2003). Data processing was performed using in-house analytic sheet under Excel (Microsoft SA, Issy-les-Moulineaux, France). Raw data, obtained from an automatic food and drink sensors (TSE, Bad Hamburg, Germany), represents for each time acquisition (a point per second) the amount of food (in mg) change. We defined a bout when there is change in the amount of food in the pellet. Each bout is then characterized by the time of its detection, its duration (in seconds) and the amount of food ingested (in mg). A meal is define as a series of bouts that are separated less than 300 seconds (intermeals intervals). A meal is then characterized by the time of its detection, its duration, the amount of food ingested, the number of bouts that composed this meals, and the average of time between two successive bouts. Meal structure are computed for a time windows of 2 hours. Results are expressed as mean ± SEM .

**Supplementary Table 1. List of primers used**

| **NPY** | **5’-** **ccgctctgcgacactacat-3’**  **5’-** **tgtctcagggctggatctct-3’** |
| --- | --- |
| **InsR** | **TqMan :** [**Mm01211875_m1**](https://www.thermofisher.com/taqman-gene-expression/product/Mm01211875_m1?CID=&ICID=&subtype=) |
| **IRS-1** | **TqMan :** [**Mm01278327_m1**](https://www.thermofisher.com/taqman-gene-expression/product/Mm01278327_m1?CID=&ICID=&subtype=) |
| **IRS-2** | **TqMan :**[**Mm03038438_m1**](https://www.thermofisher.com/taqman-gene-expression/product/Mm03038438_m1?CID=&ICID=&subtype=) |
| **GLP-1R** | **TqMan :** [**Mm00445292_m1**](https://www.thermofisher.com/taqman-gene-expression/product/Mm00445292_m1?CID=&ICID=&subtype=) |
| **PI3K** | **TqMan :** [**Mm01282781_m1**](https://www.thermofisher.com/taqman-gene-expression/product/Mm01282781_m1?CID=&ICID=&subtype=) |
| **IGF1-R** | **TqMan :** [**Mm00802831_m1**](https://www.thermofisher.com/taqman-gene-expression/product/Mm00802831_m1?CID=&ICID=&subtype=) |
| **ACTB** | **TqMan: Mm00607939_s1** |
| **R18S** | **TqMan: Hs99999901_s1** |

**Supplementary Table 2. Characteristics of the participants.**

Presented are medians [interquartile range]

| **NPY** | **Subject characteristics** |
| --- | --- |
| N | 140 (85F/55 M) |
| Age (yr) | 65 [58-71] |
| BMI (kg/m^2^) | 25.6 [23.2-27.9] |
| Weight (kg ) | 0.91 [0.85-0.96] |
| HbA1c (%) | 5.7 [5.4-5.7] |
| Plasma glucose (mg/dl) | 98 [90-108] |
| CSF glucose (mg/dl) | 61 [58-65] |
| Serum insulin (pmol/l) | 82 [53-121] |
| CSF insulin (pmol/l) | 7.7 [6.5-9.4] |
| CSF/Serum insulin | 0.095 [0.066-0.151] |
